# Finding Agreement: fMRI hyperscanning reveals dyads that explore in mental state space facilitate opinion alignment

**DOI:** 10.1101/2024.10.03.616501

**Authors:** Sebastian P.H. Speer, Haran Sened, Laetitia Mwilambwe-Tshilobo, Emily B. Falk, Lily Tsoi, Shannon M. Burns, Diana I. Tamir

## Abstract

Many prize synchrony as the ingredient that turns a discordant conversation into a delightful duet. Although failure to align is sometimes associated with discord, exploring divergent opinions can also foster understanding and agreement, satisfy curiosity, and spur imagination. Using fMRI hyperscanning and natural language processing, we tested if dyadic alignment or exploration during decision-making conversations was a more effective route to agreement. Dyads (*N* = 60) discussed pressing societal problems while being instructed to either *persuade* or *compromise* with their partner. Our analysis uncovered four key insights: First, individuals instructed to compromise rather than persuade tended to agree more at the end of the conversation. Second, hyperscanning and linguistic analyses revealed that encouraging compromise resulted in increased exploration during conversations; dyads given compromise instructions traversed more diverse mental states and topics. Third, heightened exploration was linked to greater agreement at the end of the conversation. Fourth, the effect of the compromise instructions on agreement was entirely mediated by the degree of exploration. Together, these results suggest that finding agreement may be spurred by exploration, something that happens spontaneously when people are motivated to compromise.

**Significance Statement:** Successful conflict resolution is not just about syncing perfectly—it sometimes means leaning into differences. Our study employed fMRI hyperscanning to examine if exploratory conversation strategies promote consensus. We paired participants to discuss societal issues with either a persuasion or compromise mindset. We found that dyads aiming to compromise reached more agreement because they delved deeper into each other’s minds and covered more topics; in contrast, dyads aiming to persuade their partner covered a narrower scope of topics and mental states and ended up disagreeing more as a result. The finding that exploration rather than simple alignment facilitates agreement could sharpen communication strategies in fields from politics to education and economics.

---

People do not always agree with each other. People disagree with friends, colleagues, and strangers on basic questions, such as what to eat for dinner, and on big societal questions, such as whether and how to address climate change or the cost of education. The inability to reach agreement can impede progress and have serious consequences for human well-being and social and environmental stability. What happens when people engage in decision-making conversations that ultimately reach, or fail to reach, consensus? In this study, we use fMRI hyperscanning and natural language processing to investigate what neural and linguistic dynamics support mutual agreement in a real-time conversation and how partners’ motives going into a conversation give rise to these dynamics.

## Motives to Compromise vs. Persuade Impact Dyadic Agreement

Some conversations foster agreement and harmony, while others amplify discord. A rich foundation of work on negotiation conversations suggests that social motives are important factors in determining the course of a conversation (De Dreu et al., 2000; Olekalns & Smith, 1999, 2003b, 2003a; Brett & Thompson, 2016). Dyads can bring at least two categories of motives to decision-making conversations: motives to compromise and persuade.

The motivation to persuade in a negotiation can prime a competitive mindset, wherein gains for one person are perceived as losses for the other (Beersma & De Dreu, 2002; De Dreu et al., 2001; Messick & McClintock, 1968; Pruitt & Rubin, 1986). With this competitive mindset, people often have a specific end goal in mind, leading them to direct the conversation toward a particular outcome. This focus on achieving a desired result can narrow a person’s thinking and reduce cognitive flexibility, making it harder to consider alternative viewpoints (Brynjarsdottir et al., 2012; Hommel, 2015; Zhang et al., 2020). At the same time, people are often motivated to resist persuasion to retain their sense of freedom and control (Knowles & Linn, 2004; Wheeler et al., 2007). Thus, competitive negotiators may hesitate to support and build on each others’ ideas (Decety et al., 2004; Lu et al., 2019). This typically results in a failure to identify tradeoffs that could have created more value and higher joint gains (Kong et al., 2014). Together, this work suggests that a motivation to persuade can lead to more rigid, constrained, and focused conversations and may result in lower levels of eventual agreement.

In contrast, motives to compromise can prime a cooperative mindset, characterized by prosocial intentions and a focus on maximizing joint outcomes for the self and others. Dyads with a cooperative mindset pursue a shared goal and actively process, learn, and build on each others’ ideas (Hargadon & Bechky, 2006; Vera & Crossan, 2005). As a result, the cooperative mindset has been linked to more creativity and cognitive flexibility (Bittner et al., 2016; Carmeli et al., 2015; Gelfo, 2019; Hon et al., 2014; Laureiro-Martínez & Brusoni, 2018) and an appreciation for the subjectivity of truth (Fisher et al., 2017). Cooperative motives are linked to integrative negotiation behaviors (De Dreu et al., 2000; Deutsch, 1973), which predominantly consist of sharing information about interests and opinions and then forging tradeoffs to produce high joint gains (Weingart et al., 1990). Cooperative negotiators consistently achieve higher joint gains and generate more value than competitive negotiators (De Dreu et al., 2000; Kong et al., 2014; Olekalns & Smith, 1999, 2003a, 2003b). Together, this work suggests that compromise motives can lead to more creative, flexible, and synergistic conversations and may result in higher levels of eventual agreement.

## Motives Shape Conversation Dynamics

People’s social motives influence the conversation dynamics or the strategies they use to bridge disconnects and pursue agreement. To find agreement, people might try to *align* their thoughts and feelings by honing in on key points. They might also try to *explore new ground*, offering fresh, unexpected ideas or diverging perspectives (Burns et al., 2024; Speer et al., 2024).

Psychologists and neuroscientists have prized *alignment* of mental states as a path to conflict resolution (Tunçgenç & Cohen, 2018; Valdesolo & DeSteno, 2011; Wiltermuth & Heath, 2009). Past work indicates that people agree more when they show greater linguistic (Manson et al., 2013), physiological (Behrens et al., 2020), and neural alignment (Liu et al., 2017; Wang et al., 2022; Zhang et al., 2023). Alignment may facilitate agreement because it enhances joint attention, mutual understanding, and empathy, all critical elements for cooperative behavior (Liu et al., 2017; Lu et al., 2019; Wang et al., 2022). Indeed, alignment has been associated with positive social outcomes in related contexts such as emotional support (Doré & Morris, 2018), interpersonal liking (Ireland et al., 2011; Putman & Street, 1984; Street et al., 1983), social influence (Ludwig et al., 2013), compliance (Jacob et al., 2011; Ramseyer & Tschacher, 2011) and managers-employee rapport (Balconi, 2010). However, it is also possible that alignment merely reflects shared understanding but does not create it.

*Exploring new ground,* a process by which people consider and discuss diverse opinions and topics, can foster understanding and collaboration (Lu et al., 2019; Perc & Szolnoki, 2008; Su et al., 2016; Williams et al., 2020). Although exploration of alternative views may initially appear as a divergence in mental states, dyads who more thoroughly explore may be better poised to reach eventual agreement. When individuals engage with diverse viewpoints and thoughtfully process the information, the resulting attitude change is more durable and meaningful. In contrast, when individuals think less deeply about the other person’s arguments or simply start out by agreeing, the consensus may be superficial and lack a lasting impact (Petty & Cacioppo, 1986). Entertaining diverse ideas within one’s own mind can mitigate polarized attitudes (Sassenberg & Winter, 2024). If a similar process comes into play when two minds entertain diverse ideas, this could attenuate dyadic disagreement. Exploration across minds, where people traverse a wide range of each others mental states, has been linked to beneficial social outcomes, such as more secure attachment (Nguyen et al., 2024), stronger social connection (Ravreby et al., 2022; Speer et al., 2024), improved emotion regulation (Lee & Fujita, 2023; Powers & LaBar, 2019), and more effective problem solving (Goldstone et al., 2024).

Given its association with divergent and flexible thinking, a cooperative mindset may encourage the *exploration of new ground* during a conversation. The cooperative mindset, like that associated with compromise, fosters more cognitive flexibility, creativity, and information sharing (Decety et al., 2004; Gilson & Shalley, 2004; Hargadon & Bechky, 2006; Van Knippenberg et al., 2004; Vera & Crossan, 2005). These findings may explain why motives to compromise promote more eventual agreement. In contrast, the motivation to persuade might lead to less exploration through the pursuit of winning or convincing one’s partner of a particular point. A persuader with a clear end state in mind might narrow the focus of a conversation, inhibiting the mutual sharing and processing of broad information that is useful for reaching meaningful agreement.

## The Current Study: Conversation Dynamics that Facilitate Agreement

This study aimed to identify the motives and dynamics that promote agreement in conversation. We tested the hypothesis that compromise promotes exploration, which in turn, helps people reach mutual agreement. Importantly, alignment and exploration are inherently temporal processes, defined by movement toward similar (in alignment) and novel (in exploration) mental states respectively. To examine temporal trajectories in mental states *during* conversation, we used fMRI hyperscanning (Babiloni & Astolfi, 2014; Kelsen et al., 2022; Tsoi et al., 2022)—simultaneous brain imaging of two conversation partners—as well as natural language processing of both participants’ spoken words (Jain et al., 2018). These methods, rarely used in prior work, allow us to capture the dynamics of alignment or exploration without relying on meta-cognition by participants (e.g., in post-hoc self-report) or observers (e.g., in coding recordings). We used this paradigm to test four interdependent hypotheses:

First, we investigated how social motives influence eventual agreement. Dyads engaged in two real-time conversations on how to allocate money to address two contentious issues, air pollution and the cost of tuition. They provided their opinions on how to allocate money before and after the conversations, which allowed us to measure the extent to which the conversations helped them find agreement. We experimentally induced a persuasion motive by instructing half of the dyads to get their partner to agree with them (both partners received the same instruction). We induced a compromise motive in the other half of dyads by instructing them to reach a joint decision. We expected to replicate previous findings (De Dreu et al., 2000; Kong et al., 2014; Olekalns & Smith, 1999, 2003a, 2003b), such that compromise dyads would reach higher levels of agreement than dyads instructed to persuade.

Next, we examined how the manipulated social motives shaped dyads’ conversation dynamics. Specifically, we assessed through neural and linguistic data how compromise vs. persuasion instructions shaped the extent to which dyads aligned or explored new ground. We tracked both conversation partners’ mental states using a brain and linguistic state decoding approach we developed previously (Speer et al., 2024). Prior research shows that three key dimensions—social impact, rationality, and valence (the “3D mind model”; Tamir et al., 2016; Tamir & Thornton, 2018; Thornton & Tamir, 2020)—capture how the brain represents underlying mental states. A person’s position within this three-dimensional framework reveals much about their internal state, and observing how two individuals move relative to each other in this space reveals their interpersonal dynamics. In particular, exploratory dyads will move farther apart (rather than closer together) within the space and cover more diverse mental states. The conversation content also offers us a window into whether dyads explore diverse ground. We used topic modeling to assess what dyads talked about throughout their conversations. Dyads could evince exploration by covering more topics, having more topic switches, and taking more distant topic jumps. We measured each of these types of topic exploration here. We tested if compromise dyads explored more than dyads instructed to persuade.

Third, we tested whether these conversation dynamics also predicted agreement—that is, whether dyads who engaged in more mental state and topic exploration agreed more by the end of the conversation.

Finally, we tested whether the conversation dynamics mediated the effect of motives on agreement. To accomplish this, we conducted a mediation analysis to assess whether the instructions to compromise led to greater agreement by encouraging greater exploration. We implemented this multi-modal approach in combination with naturalistic free-flowing conversations to be able to capture the emergent properties of social interactions.

## Results

### The Instruction to Compromise Amplifies the Positive Effect of Conversation on Agreement

We first tested how social motives (*compromise vs. persuade*) influence eventual agreement. Prior to each conversation, participants were exposed to a discussion problem and five different solutions to address the problem. Participants independently allocated a hypothetical 100 million dollars to five different solutions both pre- and post-conversation, which allowed us to measure how similar dyads’ opinions were before and after the conversation. Participants agreed significantly more after the conversations than before (Figure 1: *t(216)* = 6.96, two-tailed, *d* = 1.10, *p* < 0.001). Increased agreement was observed in both conditions (compromise: *t(108)* = 6.36, two-tailed, *d* = 1.42, *p* < 0.001; persuade: *t(106)* = 3.51, two-tailed, *d* = 0.79, *p* < 0.001). Next, we tested how compromise vs. persuade motives influence this relationship. We found a significant interaction effect between time and condition (Figure 1: *t(216)* = 2.13, two-tailed, *d* = 0.34, *p* =0.03), such that instruction to compromise amplified this effect: participants in the compromise condition agreed significantly more than participants in the persuade condition after the conversation (Figure 1: *t(54)* = 3.28, two-tailed, *d* = 0.89, *p* = 0.002) but not before the conversation (Figure 1: *t(54)* = 0.84, two-tailed, *d* = 0.23, *p* = 0.40). Thus, talking to each other and exchanging perspectives appears to increase agreement, and being instructed to compromise enhances this effect.

**Figure 1.**
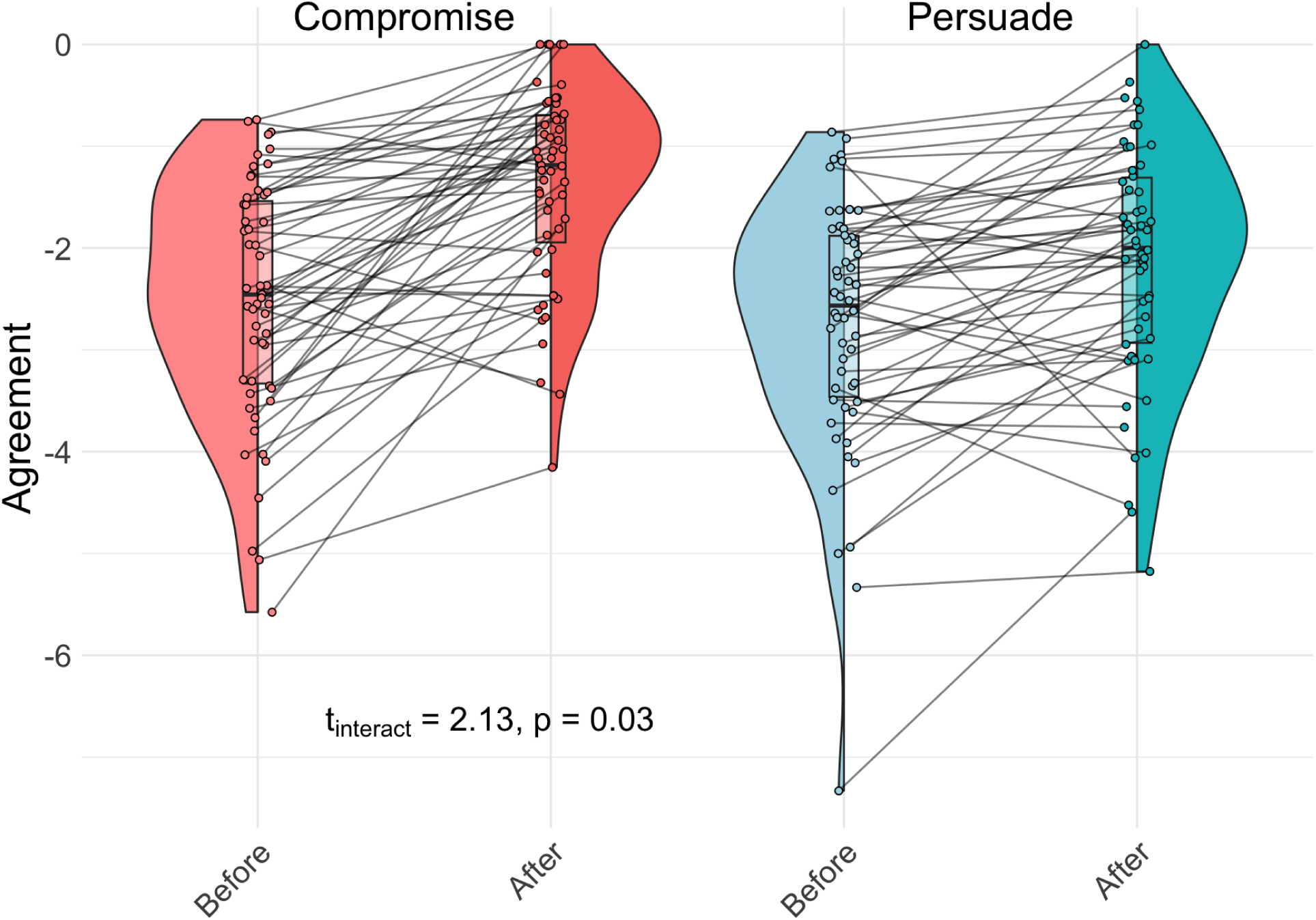
Participants agree significantly more after vs. before the conversation, with this effect amplified in the compromise condition. Agreement represents the inverse distance in opinion space (agreement = −1* distance such that higher values reflect higher agreement).

### Instruction to Compromise Increases Exploration

Social motives matter: being instructed to compromise increased the amount that conversation partners agreed with each other. Next, we investigated if instructions to compromise also increased exploration during the conversation. We quantified exploration by tracking how dyads moved through mental state and topic space. To measure how dyads moved through mental state space, we located each partner on three mental state dimensions throughout the conversation. We located dyads in neural mental state space by applying a neural decoding model to both partners whole-brain neural activity at each moment of the conversation; we located dyads in linguistic mental state space by using natural language processing on the language each speaker produced over the course of the conversation (Speer et al., 2024; Supplementary Material 1). To measure how dyads moved through topic space, we applied unsupervised machine learning to extract the topic of each turn and calculate the semantic distance between adjacent topics (Supplementary Material 2).

#### Mental State Exploration

Using the decoded neural coordinates, we measured mental state exploration as the increase in distance between participants over time (divergence) and the diversity of mental states covered by the dyad collectively (novelty). Greater divergence over time and greater novelty represented more exploration. We expected the instruction to compromise to induce exploration and the instruction to persuade to reduce exploration. We employed multilevel models to examine if dyads diverged over the course of a conversation. There was a two-way interaction between time and social motive, such that dyads instructed to compromise diverged in neural mental state space, whereas dyads instructed to persuade converged (*β_std_* = −0.06, 95% CI [-0.0007, −0.0003], *SE* = 0.00008, *p* < 0.0001; Figure 2A). Dyads motivated to compromise were aligned more initially than dyads motivated to persuade and then diverged in mental state space until their distance was larger than the persuaders’. The linguistic analysis replicated this interaction effect (*β_std_* = −0.22, 95% CI [-0.08, −0.02], *SE* = 0.016, *p* = 0.001; Figure 2B), such that dyads in the compromise condition diverged in mental state space, whereas participants in the persuade condition converged.

**Figure 2.**
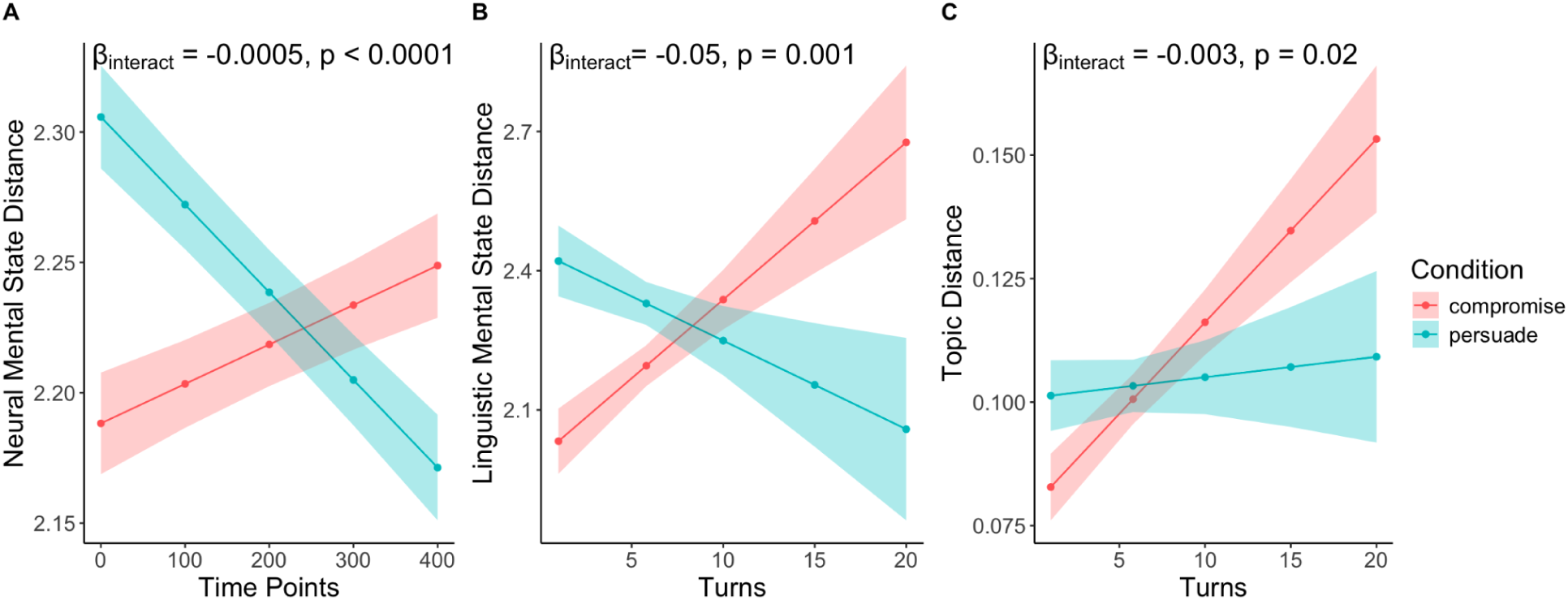
The instruction to compromise increases exploration. Compromisers diverge while persuaders align in neural (A) and linguistic mental state space (B) and in topic space (C). We used time points (TRs) and turns, respectively, as the smallest unit of analysis for the neural and linguistic data.

To measure dyadic novelty, we calculated the difference between the distributions of the mental state locations for each turn to all previous turns (see Methods). We measured speaker and listener mental states using the simultaneously collected neural data and sequential speaker states in the linguistic data. We expected the compromise condition to be associated with more novelty. The neural novelty analysis revealed a trend suggesting that dyads in the compromise condition exhibited more novel mental states than participants in the persuade condition (*t(53)* = 1.95, two-tailed, *d* = 0.53, *p* = 0.056; Figure 3A). The linguistic novelty analyses revealed a significant effect in the same direction, such that dyads in the compromise condition exhibited significantly more novel mental states than dyads in the persuade condition (*t(53)* = 3.19, two-tailed, *d* = 0.88, *p* = 0.002; Figure 3B).

**Figure 3.**
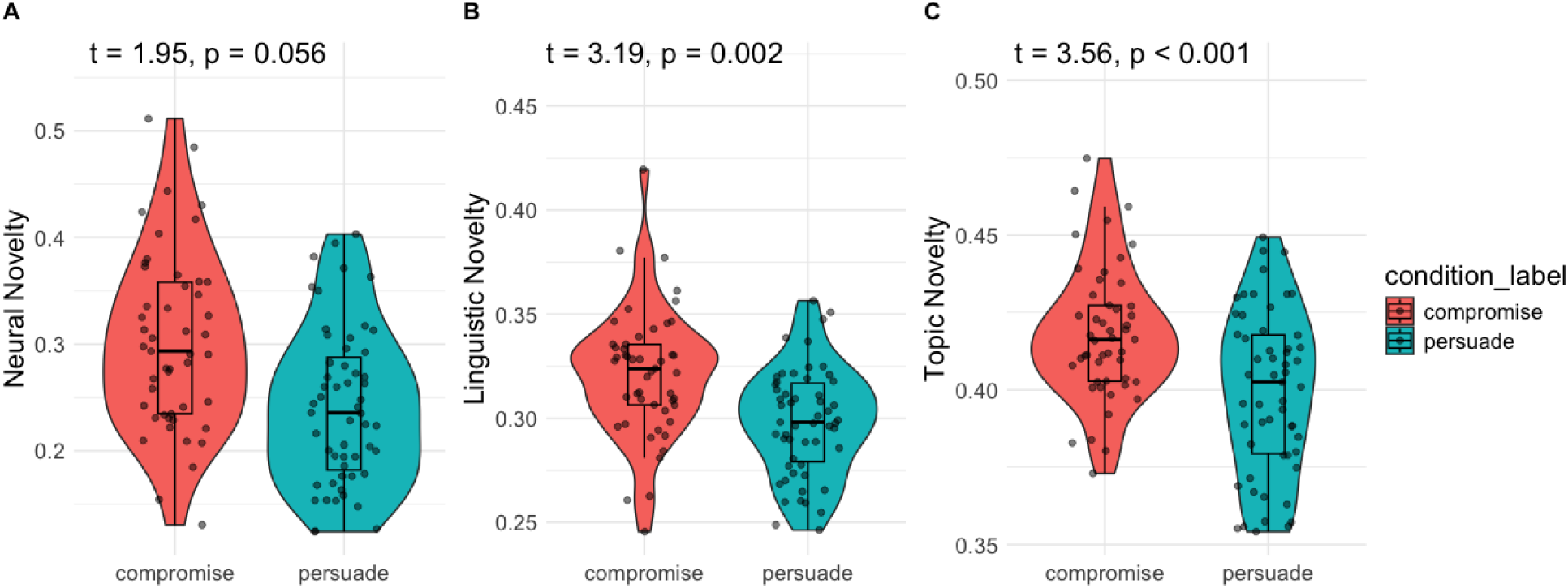
Dyads instructed to compromise exhibit more novelty in their mental state and topic trajectories. (A-B) Dyads in the compromise condition have more diverse mental state trajectories than dyads in the persuade condition and (C) jump longer distances on average between topics.

Together, these findings suggest that the instruction to compromise increases mental state exploration both in terms of change in relative distance (divergence) and absolute novelty.

#### Topic Exploration

To measure topic exploration, we applied unsupervised machine learning to extract the topic of each turn and calculate the semantic distance between them (Supplementary Material 2). We expected dyads in the compromise condition to explore more topic space, which we measured in several ways. To parallel the mental state analyses, we first looked at divergence in topic space between conversation partners. This analysis revealed a two-way interaction effect between turns and social motives on topic distance: Dyads instructed to compromise explored increasingly more distinct topics than dyads instructed to persuade (*β_std_* = −0.15, 95% CI [-0.006, −0.0005], *SE* = 0.001, *p* = 0.02; Figure 2C). Thus, similar to the mental state findings, dyads instructed to compromise diverge in topic space over time, whereas dyads instructed to persuade do not.

We also found that dyads in the compromise explored more novel topics, as evidenced by longer distances between topics (*t*(53) = 3.56, two-tailed, *d* = 0.98, *p* = 0.0008; Figure 3C), mirroring the mental state novelty analyses. In addition, compromise dyads generated significantly more topics (*t(53)* = 2.61, two-tailed, *d* = 0.72, *p* = 0.01) and showed a trend towards switching topics more frequently (*t(53)* = 1.99, two-tailed, *d* = 0.55, *p* = 0.051).

Collectively, these findings demonstrate that dyads instructed to compromise explore a more expansive topic space by switching between more distant topics, whereas conversationalists motivated to persuade stick with similar topics for longer.

### Exploration is Associated with More Agreement

After seeing that instructions to compromise causally increased both agreement and exploration, we next tested if exploration enhances agreement. We expected that exploration through both mental state and topic space would result in more agreement.

#### Mental State Exploration

We fit two multilevel regression models (neural and linguistic) to investigate how agreement affects divergence over time between partners in mental state space. Each model predicted distance from agreement and time points (within-dyads, for neural) or turns (within-dyads, for linguistic) and controlled for initial difference in opinion (distance in opinion space before the conversation). The analyses revealed significant interactions between time points and agreement, such that dyads who agreed more diverged more in neural mental state space (β_std_ = 0.01, SE = 0.00004, 95% CI [0.00004, 0.0002], p = 0.003; Figure 4A). Dyads who agreed more started closer in mental state space than dyads who disagreed more and then diverged in neural mental state space until their distance was larger than the dyads who disagreed more. No significant effects were found for the effect of agreement on linguistic mental state exploration.

**Figure 4.**
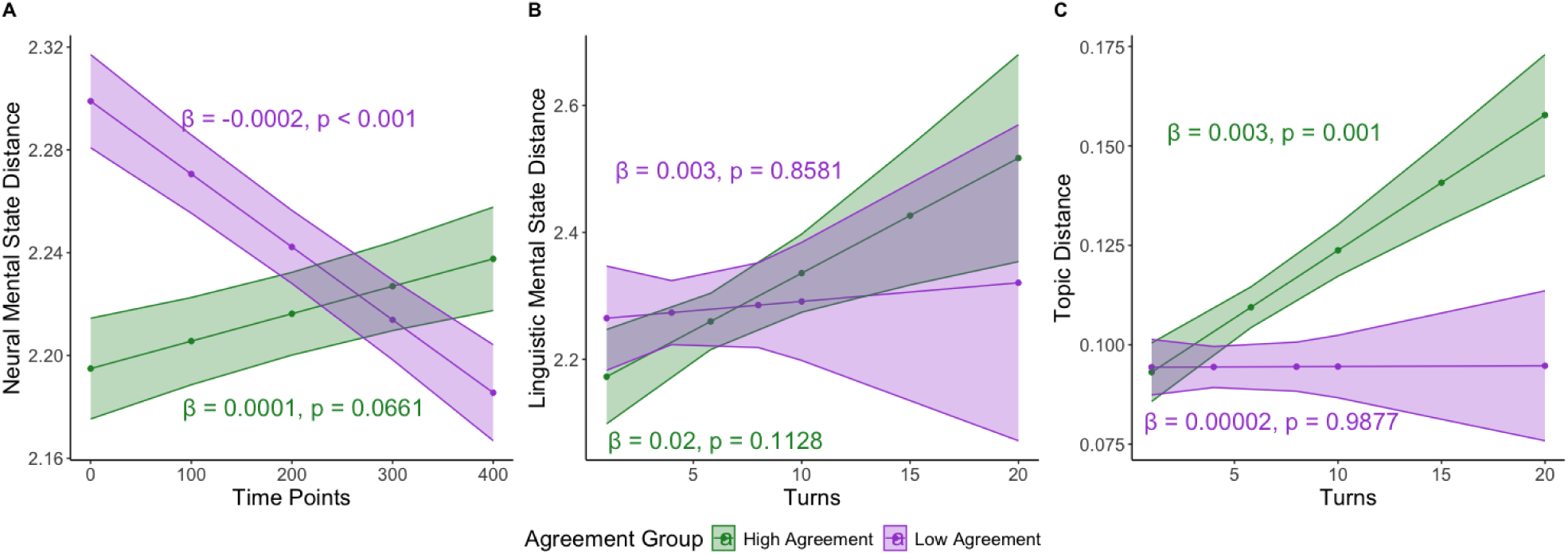
Dyads who agreed more diverged more in mental state and topic space. These plots show the fitted values of two separate models fit for the data median split on agreement. The βs represent the slopes for distance over time (timepoints/turms) for the high and low agreement groups.

We also found that agreement was significantly associated with both neural mental state novelty (β_std_ = 0.30, 95% CI [0.62, 7.55], SE = 0.1.72, p = 0.02; Figure 5A) as well as linguistic mental state novelty (β_std_ = 0.29, 95% CI [4.39, 17.33], SE = 3.21, p = 0.002; Figure 5B).

**Figure 5.**
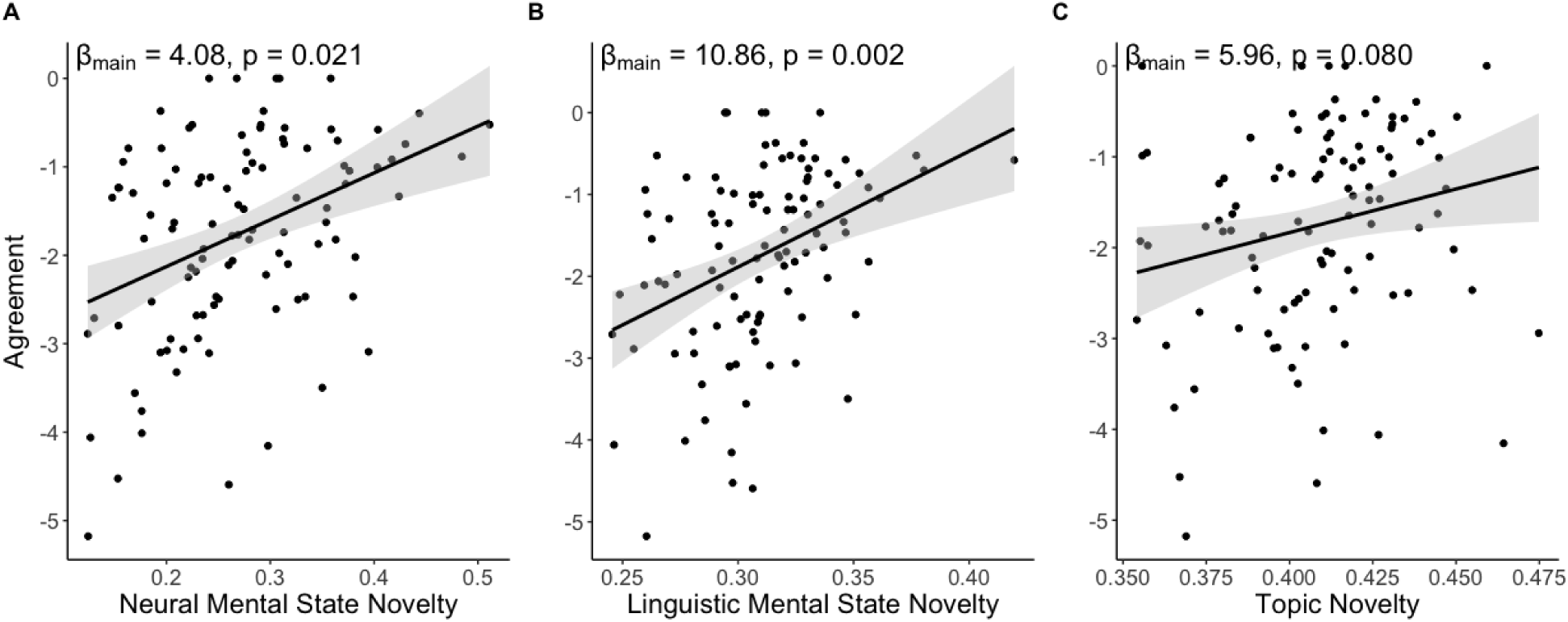
Dyads who exhibit more novelty in their topic and mental state trajectories agree more. The grey ribbons represent the 95% confidence intervals around the fitted values.

Together, this suggests that more mental state exploration leads to more agreement.

#### Topic Exploration

Paralleling the mental state analyses, we fit a multilevel regression model to investigate how agreement relates to changes in distance in topic space over time. We found that dyads who agreed more diverged more in topic space over time, whereas dyads who disagreed more did not diverge (β_std_ = 0.07, 95% CI [0.00005, 0.003], SE = 0.0006, p = 0.042; Figure 4C).

We also found that agreement was marginally associated with topic novelty (β = 5.96, β_std_ = 0.14, 95% CI [-0.74, 12.66], SE = 3.33, p = 0.08; Figure 5C), similar to the mental state novelty findings. In addition, dyads who generated more topics (β_std_ = 0.17, 95% CI [0.002, 0.059], SE = 0.01, p = 0.039) found more agreement. No significant effects were found for topic switches. In sum, these findings imply that more topic exploration leads to more agreement.

### Exploration Mediates the Effect of Social Motive on Agreement

So far, we have shown that being instructed to compromise increased both agreement and exploration. Exploration, in turn, increased agreement. Here, we tested whether the effect of social motives (compromise vs. persuade) on agreement was mediated by exploration. To this end, we created a composite measure of exploration by computing the mean over the neural mental state, linguistic mental state, and topic novelty. A mediation analysis revealed that social motive significantly predicted exploration (*β_std_* = 0.32, 95% CI [0.009, 0.04], *SE* = 0.008, *p* = 0.003; Figure 6), and exploration significantly predicted agreement (*β_std_* = 0.27, 95% CI [1.88, 13.68], *SE* = 2.92, *p* = 0.011; Figure 6). After controlling for exploration, the direct effect of social motive on agreement was no longer significant (*β_std_* = 0.14, 95% CI [-0.08, 0.73], *p* = 0.12; Figure 6), indicating full mediation. The indirect effect of social motive through exploration was significant (*β_std_* = 0.09, 95% CI [0.02, 0.18], *p* = 0.008; Figure 6) based on 5000 bootstrap resamples. The total effect of social motive on agreement was also significant (*β_std_* = 0.23, 95% CI [0.05, 0.41], *p* = 0.004; Figure 6). These findings show that the effect of instruction to compromise on agreement is fully mediated by exploration.

**Figure 6.**
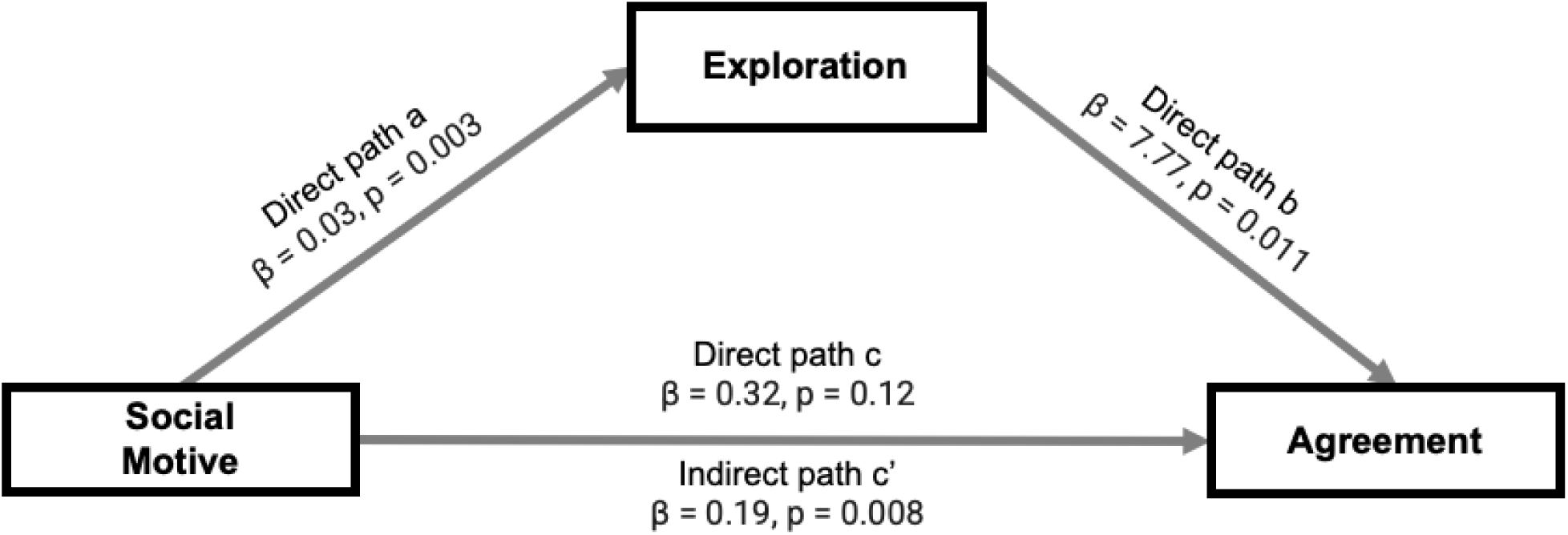
Composite measure of exploration mediates the positive effect of the instruction to compromise on agreement.

These results were robust across several analytic decisions. First, we used a composite across the three measures because analyses revealed that all three modalities (neural, linguistic, and topic) tap into the same underlying construct (see Supplementary Material 3) and because the composite measure best predicts agreement (see Supplementary Material 4). That said, mediation analyses with each individual measure of exploration replicated this same pattern of results (Supplementary Material 5). Second, we used the novelty measure because it offers a more intuitively interpretable metric (i.e., absolute exploration) than divergence over time (see Discussion) and because it was found to be more predictive of agreement at the end of the conversation (see Supplementary Material 6). That said, mediation analyses using the divergence measure did not show a significant mediation effect (see Supplementary Materials 6).

## Discussion

The ability to find agreement with others can facilitate a wide range of human goals and achievements. How do people effectively navigate conversations to reach this end? In this study, we used fMRI hyperscanning and linguistic analyses to investigate how people use exploration as they discuss pressing societal problems to reach agreement. Our analyses yielded four insights that collectively guide us toward an answer. First, people instructed to compromise as opposed to persuade agreed more at the end of the conversation. Second, compromise increased mental and topic exploration. Third, exploration was associated with more agreement. Finally, the effect of compromise on agreement was fully mediated by the extent of this exploration. Thus, compromising may facilitate agreement by promoting greater exploration during conversations.

Our study found that people instructed to compromise explore more mental states and topics than those instructed to persuade. A compromise motive may lead to more exploration for at least two reasons. First, social motives can shape the conversation dynamics (Brett, 2000; Brett & Thompson, 2016). Compromise motives may lead to more exploration by fostering open dialogue and information sharing (De Dreu et al., 2000; Deutsch, 1973; Weingart et al., 1990), which may induce more diverging mental states manifested in more dissimilar neural patterns. Second, dyads instructed to compromise start more aligned at the beginning of the conversation, whereas dyads instructed to persuade start off more distant in mental state space (see Figure 2). By entering the conversation already ‘on the same page’ in terms of their motivational stance and mental states, compromise dyads can more safely engage divergent interests and perspectives. Here, the more dyads initially agreed on solutions, the more they diverged in conversation (See Supplementary Material 6). Dyads who do not arrive already on common ground may need to initially focus on establishing common ground; only once they reach a critical threshold of alignment can they start diverging (Sened et al., 2024). It is important to note that this initial gap in mental state alignment is not due to differences in participants’ starting opinions, which were similar across conditions (see Figure 1). Initial agreement and mental state alignment independently contribute to facilitating exploration.

Our study also revealed that exploration leads to more agreement. Exploring new territory brings novelty and unpredictability to discussions, making them more engaging, meaningful, and productive. People often look for novel topics (Mehl et al., 2010), and when conversations feature this level of depth and surprise, they tend to foster greater enjoyment (Cooney et al., 2017; Reece et al., 2022), boost overall well-being (Mehl et al., 2010), reduce negative emotions (Slepian & Moulton-Tetlock, 2019), strengthen social connections (Aron et al., 1997; Speer et al., 2024), and alleviate boredom (Westgate & Wilson, 2018). In a negotiation context, exploration can boost conflict resolution by widening the range of possible solutions (Hoehl et al., 2021; Koban et al., 2019) allowing for more original and unpredictable approaches (Dideriksen et al., 2023; Speer et al., 2024), and preventing suboptimal outcomes (Abney et al., 2015; Shirado & Christakis, 2017; Wallot et al., 2016). An exploratory approach, characterized by open questions and active listening, builds trust and creates a safe space (Brooks et al., 2015; Jones, 2011; Yeomans et al., 2019). This approach allows partners to learn about each other’s negotiation interests, motives, and perspectives (Fisher et al., 2011; Walton & McKersie, 1965), generating novel and creative insights (Bittner et al., 2016; Carmeli et al., 2015; Gelfo, 2019; Hon et al., 2014; Laureiro-Martínez & Brusoni, 2018). Exploration may also foster psychological distance, which in turn promotes abstract thinking and shifts attention toward broader, essential features rather than minor details (Liberman & Trope, 2014; Trope et al., 2007).

Crucially, our final key finding shows that the effect of compromise on agreement is fully mediated by exploration. This aligns with past research (De Dreu et al., 2001; Olekalns & Smith, 1999, 2003a, 2003b; Weingart et al., 2007) suggesting that the motivation to compromise fosters greater consensus. Here, we offer a mechanism for how this happens, suggesting that compromise motives enhance agreement by amplifying exploratory dynamics. Motives thus set the stage for emergent conversation processes that ultimately shape social outcomes: even when parties share compatible motives, ineffective dynamics can lead to misunderstandings and stall progress, while effective dynamics can bridge divides despite initial misalignment. Consequently, future research and practice in conversation and joint decision-making may benefit from interventions that strengthen exploratory dynamics to improve outcomes.

These findings on the benefits of exploration for social outcomes may seem at odds with previous research showing that aligning movements, language and neural states increases agreement (Behrens et al., 2020; Krueger et al., 2007; Shaw et al., 2018; Sievers & Thornton, 2024; Tunçgenç & Cohen, 2018; Valdesolo & DeSteno, 2011; Wiltermuth & Heath, 2009). Why might alignment promote cooperation in some cases and exploration in others? First, previous fMRI hyperscanning research on synchrony has focused on highly controlled, non-verbal decision-making tasks, in which participants—without conversing—took turns making financial choices under strict timing and stimulus presentation constraints (such as the Prisoners Dilemma or the Trust Game; e.g. Behrens et al., 2020; King-Casas et al., 2005; Krueger et al., 2007; Manson et al., 2013; Shaw et al., 2018). By scanning people during naturalistic conversations, our paradigm introduces linguistic nuances, emotional cues, and dynamic turn-taking—all of which may add complexity and enable exploration that may have been unavailable in purely non-verbal contexts. Second, prior studies have largely focused on movement synchrony, physiological alignment (e.g., heart rate, skin conductance), or speech entrainment—all relatively direct and lower-dimensional metrics associated with cooperation. Even when studies measure neural alignment, this is primarily captured as inter-subject correlation across averaged brain activity in specific spatially consistent regions. In contrast, our measures of neural alignment capture a more complex, high-dimensional signal, which may be the space in which exploration can be uniquely revealed and where it may matter most for social outcomes. Specifically, we measured coordinated trajectories through a ‘mental state space’—capturing the concepts individuals represent during conversation. This shift is more than a methodological pivot; it underscores how people can attend to the same things yet differ substantially in their underlying mental content. Third, context matters: previous studies span a range of contexts, and some explicitly compare these different settings to examine how they shape the interaction dynamics and outcomes. While the findings in the literature are mixed, the majority of studies suggests that synchrony is higher in affiliative rather than task-based conversations (Dideriksen et al., 2023) and in competitive rather than cooperative contexts (Liu et al., 2015, 2017; Wang et al., 2022). Therefore, future studies that systematically compare different tasks, measurement modalities, and contexts will be critical to disentangling how these factors interact to shape the underlying interaction dynamics that promote cooperation.

Is it enough for just one person to be exploratory, or does it require both partners? Our findings suggest that while both matter, *dyadic* exploration is more strongly associated with increased agreement than individual exploration alone (see Supplementary Material 4). This extends previous studies (Kong et al., 2014) by emphasizing the emergent properties of coordinated exploration rather than one person pulling the other along. Reciprocity may be key here (Gouldner, 1960): agreement emerges when both partners share information and build on each other’s ideas (Liu & Wilson, 2011). Dyadic exploration may help partners overcome conversational impasses and break free from stagnation (Druckman, 1986; Putnam & Jones, 1982; Weingart et al., 1990). Negotiation research shows that impasses can be disrupted when partners recognize stalled progress and decide to share more openly (Brett et al., 1998; Druckman, 1986). To understand the determinants of agreement, we must study how partners dynamically explore and shape each other’s opinions in natural conversations; it’s the interplay between both participants—the shared journey through ideas and mental states—that fosters genuine consensus.

These findings were observed across multiple convergent modalities and measures. First, we tracked exploration through two experiential and conceptual spaces essential for dyadic conversation: mental states and topics. Mental state space captures the mental content of both the self and others; the ability to navigate this space through ‘mentalizing’ is crucial for successful social interactions. Topic space captures the linguistic content that dyads cover. Tracking motion through this space provides insights into how many diverse aspects of a problem dyads consider, revealing the creativity and flexibility of their problem-solving approach. Second, we measured exploration using both neural and linguistic data. By combining the high resolution of neural measures with the scalability of linguistic measures, we achieve a comprehensive and efficient understanding of exploratory dynamics. Finally, we operationalized exploration in terms of divergence and novelty. We directly tested convergent validity across all our metrics and modalities, finding significant positive correlations (see Supplementary Material 3). This convergent evidence across diverse modalities and dimensions enhances the validity of our conclusion that exploration facilitates agreement. Such multi-measure corroboration underscores the promise of exploratory dynamics as a target for interventions.

That said, our two measures of exploration, divergence and novelty, captured different aspects of the conversation. That is, even though the two measures of exploration are correlated (see Supplementary Material 3), their differences may be informative. First, only novelty predicted agreement at the end of the conversation (see Supplementary Material 6). As a result, only novelty emerged as a significant mediator of the effect of compromise on agreement (see Supplementary Material 5). Second, divergence but not novelty was associated with initial agreement (see Supplementary Material 7). Why might these correlated measures show different profiles? Novelty captures the absolute amount of space covered, whereas divergence captures the change in relative distance between the two conversation partners. Two conversation partners can remain at a significant relative distance from each other (high divergence) without extensively exploring new areas of the conversational space (low novelty), as they may hold different positions within familiar territory without venturing into new topics or ideas. Because novelty tracks how much new ground the partners collectively explore, it is closely tied to whether they discover common ideas that help bridge their differences. Divergence, in contrast, measures how far apart partners remain, regardless of whether they introduce genuinely new topics. Early in the conversation, having initial agreement can create a sense of security that allows partners to feel comfortable with diverging in mental state space—moving far away from each other’s positions—and this divergence itself can be a prerequisite for novelty, as it frees them to explore new and uncharted territories. Consequently, novelty aligns more directly with processes that foster agreement through the discovery of common ideas, while divergence reflects the distance between participants without necessarily indicating progress toward a shared understanding. Therefore, it may not be enough to just diverge from each in mental states and topics; coordinated exploration of truly novel mental and topic territory may be required to find agreement.

This study focused on comparing the effectiveness of compromise and persuade motives in shaping the neural and conversational dynamics that promote agreement. This work sets the stage for several promising avenues for future research. First, it is important to explore what differentiates successful persuaders from unsuccessful ones. Within the persuade conditions, there was significant variance in how well individuals convinced their partner of the merits of their initial opinions. What strategies grant a persuader the power to pull the other towards themselves in opinion space? In our study, persuaders were pitted against persuaders, making effective persuasion all the more impressive. Second, future studies may benefit from exploring how conversation dynamics change when partners arrive with different motives. In particular, conflictual conversations would amplify the potential impact of effective strategies. For example, participants could arrive with more polarized opinions from the start, or dyads’ personal incentives could be pitted against each other so that gains for one mean direct losses for the other. How can such dyads bridge such a divide in initial opinions and motives, and would exploration still offer a route toward ultimate agreement? Finally, it will be essential to determine whether our results hold when the personal and global stakes are higher and more real than the conditions we constructed in the lab. Our findings align with prior work suggesting that training employees, politicians, and teachers in integrative negotiation and communication skills can have positive effects on bridging divides (Lewicki et al., 2024; Thompson & DeHarpport, 1994), and we suggest that future research that bridges the current findings with these broader contexts would be fruitful.

This study addresses a key question in conversation science: What leads to mutual agreement in conversations? The findings show that overall conversations increase agreement. However, not all conversations are equally conducive to the alignment of opinions: conversations that are aimed at finding compromise as opposed to persuading lead to more agreement. This effect can fully be explained by the fact that compromise bolsters exploration in conversation, which in turn facilitates agreement. Compromise-oriented conversations were characterized by more novel mental state trajectories and more diverse and distant topics. These insights inform broader debates on how to reach agreement and provide useful strategies for navigating disagreement constructively.

## Methods

The primary aim of the study was to investigate how people navigate conversations to achieve mutual agreement. To this end, we used fMRI hyperscanning: 60 dyads engaged in real-time conversations on discussion problems aimed at stimulating diverging opinions. Half of the recruited dyads were instructed to compromise, whereas the other half were instructed to persuade. We first tested how social motives (compromise vs. persuade) relate to agreement. Secondly, we examined how social motives influence conversation dynamics (*alignment* vs. *exploration*). Third, we assessed how conversation dynamics influence agreement. Lastly, we assessed whether conversation dynamics mediate the effect of motives on agreement.

### Participants

A total of 60 dyads (120 participants) engaged in a real-time conversation while both dyad members were simultaneously scanned using fMRI hyperscanning. Due to technical issues, the data for four dyads remained incomplete. The final dataset consisted of 56 dyads, which were randomly assigned to compromise (29 dyads; *n* = 58 participants; age 17 - 36, *M*_age_= 19.94, *SD_age_*= 3.05; 31 women, 27 men, 0 non-binary, African American = 1, Asian = 12, Caucasian = 28, Other = 17), or to persuade (27 dyads; *n* = 54 participants; age 18-23, *M_age_* = 19.66, *SD_age_* = 1.27; 38 women, 15 men, 1 non-binary, African American = 6, Asian = 17, Caucasian = 22, Other = 9). All dyads were randomly paired and were unacquainted before the study. All participants had to be at least 18 years old to be eligible for the study and had to be cleared for undergoing an MRI scan (e.g. no metal implants, heart valves etc.).

### Protocol

Participants were simultaneously scanned in two MRI scanners while they learned about and discussed two discussion problems (air pollution and cost of tuition). These discussion problems were chosen based on behavioral pretesting on what current societal problems inspire the most diverse opinions regarding solutions.

Participants were first introduced to one of the discussion problems by reading a summary of the problem (see Figure 7). Next, participants were presented with five different potential solutions to address the problem they had just learned about. Participants were then asked how $100 million should be allocated across the five options to address the discussion problem. They could take as much time as they wanted to answer the questions. Afterward, they listened to three news clips about the discussion problem for about seven minutes. The news articles were chosen for their descriptive reporting of people experiencing consequences of the public health problem, rather than promoting any particular solution to the problem, in order to minimize influence on participant opinions.

**Figure 7.**
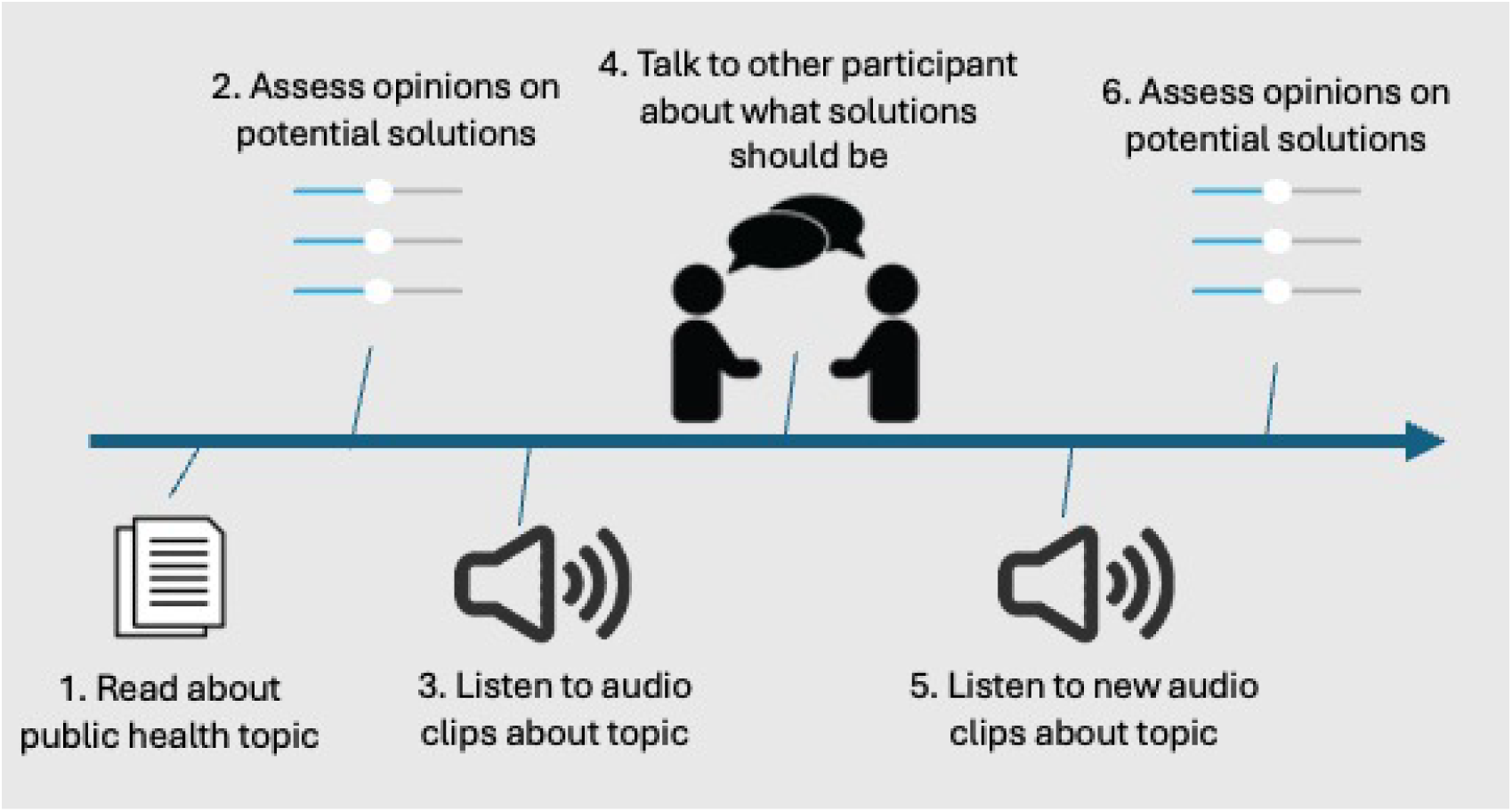
The order of the tasks. Participants first read about the public health problem and then provided their opinions about five potential solutions. Afterwards, they listen to an audio clip about the discussion problem. Next, they discussed the problem with their conversation partner before providing their opinion about the solutions a second time. Lastly, they evaluated the conversations.

Just before starting the first conversation, dyads were assigned to one of two between-dyad conditions: (i) Compromise: Dyads were asked to find a compromise on their opinions on the discussion problems presented. (ii) Persuade: Dyads were instructed to persuade their conversation partner of their opinion. Both participants received the same instruction in each condition for both conversations (i.e. both were instructed to compromise or both were instructed to persuade). We chose a between-subject design because it would be hard for dyads to switch goals (persuasion vs. compromise) from one trial to the next. Participants then conversed with their partner for 10 minutes. The conversation portion of the trials began with the conversation prompt stating, “Please discuss the solutions with your partners” (3 sec), followed by 10 min of turn-taking (see Figure 8). The conversations were freeform, which meant that participants could say whatever they wanted. Participants indicated the end of a turn by pressing a button to transfer the speaker role to the conversation partner.

**Figure 8.**
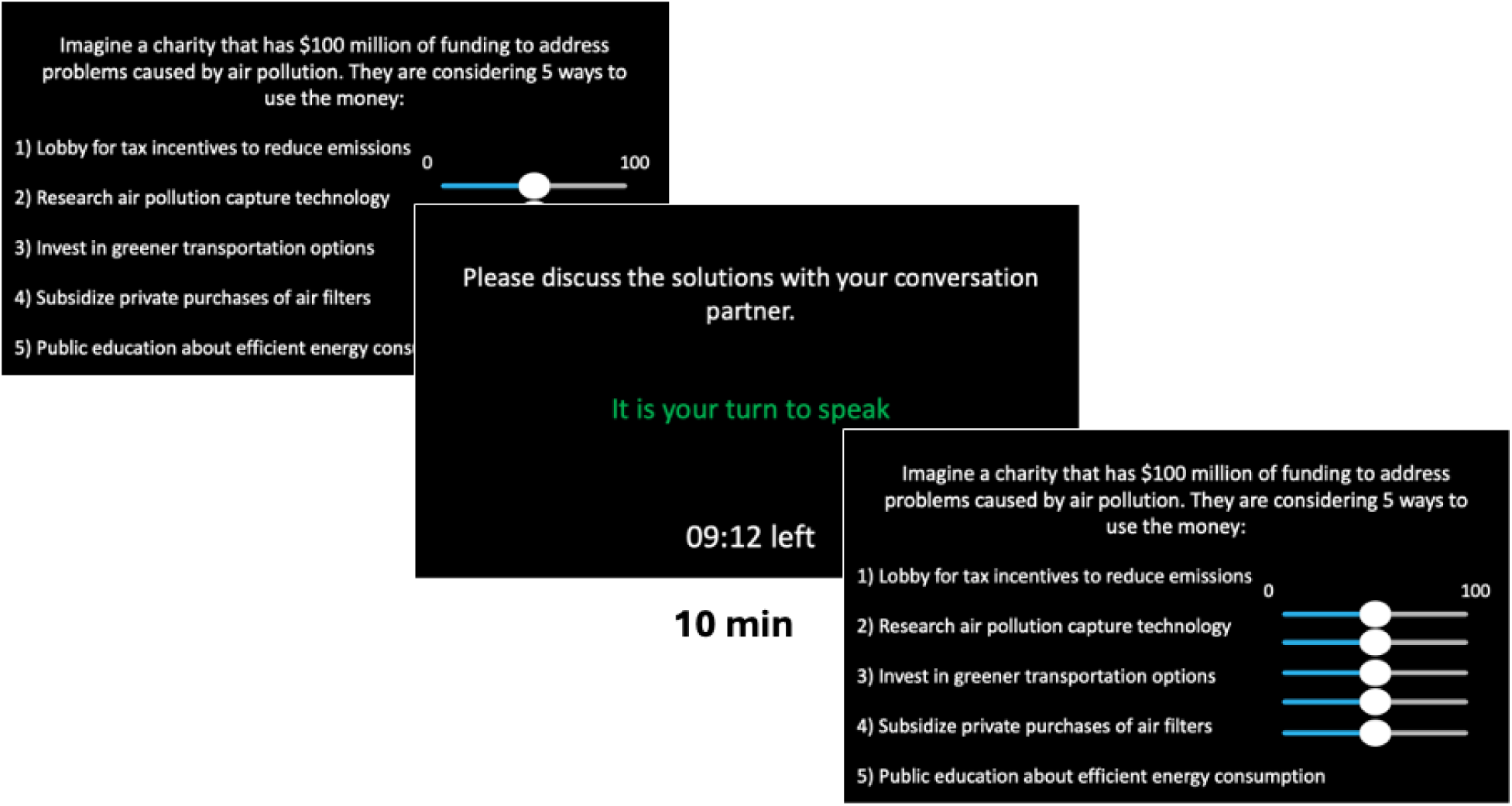
An example trial as seen from a participant’s perspective. Dyads viewed the discussion prompt and allocated $100,000,000 across the five proposed solutions for addressing the problem. Subsequently, they were assigned speaker and listener roles, and the speaker started speaking. The first speaker pressed a button to ‘pass the mic’ to the listener, who then became the speaker. Participants saw how much time there was left in the current trial at the bottom of the screen. After they finished the conversation, they were asked to reallocate money across the five different solutions.

After participants finished the conversation, they again watched about seven minutes of three news clips (different from the first viewing) and again answered how they would allocate $100 million to five different solutions addressing the problem.

After completing all parts of the task for the first discussion problem, participants repeated all parts for the second discussion problem. The discussion problem they discussed first was counterbalanced across dyads.

### Speech Recording

We recorded all conversations during the fMRI scan using a specialized MR-compatible recording system (FOMRI II; Optoacoustics Ltd). This system employs two optical microphones positioned orthogonally. One microphone captures background noise (reference), while the other captures both background noise and the speaker’s speech (signal). A dual-adaptive filter subtracts the reference input from the signal channel using a least mean square method. To ensure effective subtraction, the reference signal undergoes adaptive filtering, with the filter gains continuously adjusted based on the residual signal and reference input. To prevent filter divergence during speech, a voice activity detector is incorporated into the algorithm. Additionally, a speech enhancement spectral filtering algorithm further preprocesses the speech output for real-time enhancement. Participants were able to communicate via an audio link, connecting the two adjacent rooms.

### Imaging Procedure

The fMRI scans for the dyads were conducted using both a 3T Siemens Skyra MRI and a 3T Siemens Prisma MRI located in adjacent rooms, with identical scanning parameters employed for both scanners. Functional scans were obtained covering the entire brain in an interleaved manner, with a slice thickness of 3.0 mm, an in-plane resolution of 3.0 × 3.0 mm, and a flip angle of 80°. The echo time (TE) was set at 28 ms, and the repetition time (TR) was 1500 ms. Additionally, a T1-weighted image was obtained to serve as an anatomical reference, with a resolution of 1.0 × 1.0 × 1.0 mm, 176 sagittal slices, a flip angle of 9°, TE of 2.98 ms, and TR of 2300 ms. To minimize head movement during scanning, subjects’ heads were secured using foam padding.

The fMRI data were preprocessed using fMRIPrep version 20.2.0, a Nipype-based tool chosen for its robustness and reproducibility in preprocessing diverse datasets. fMRIPrep automatically adapts workflows based on best-in-class algorithms, enabling high-quality preprocessing with minimal manual intervention. Detailed methods for preprocessing can be found in (Speer et al., 2024).

Each T1-weighted (T1w) volume was corrected for intensity nonuniformity and skull stripped. Spatial normalization was performed using the International Consortium for Brain Mapping (ICBM) 152 Nonlinear Asymmetrical template version 2009c through nonlinear registration. Brain tissue segmentation of cerebrospinal fluid (CSF), white matter (WM), and gray matter was carried out on the brain-extracted T1w. Field map distortion correction involved coregistering the functional image to the same-subject T1w image, followed by boundary-based registration to the T1w with nine degrees of freedom. Motion correction, field distortion correction, and spatial normalization were applied in a single step using Lanczos interpolation. Physiological noise regressors were extracted using CompCor, with six components each for temporal (tCompCor) and anatomical (aCompCor) variants.

Because speaking induces motion artifacts and potential distortions in the magnetic field, an additional denoising step was necessary to clean the data after preprocessing. Consequently, we conducted further confound regression on the preprocessed data. This regression involved incorporating various regressors, including six head motion parameters, their cosine, first derivatives, squares, signals from white matter and cerebrospinal fluid, along with their derivatives and squared derivatives, as regressors of no interest. The residuals obtained from this regression, which represent the cleaned data, were then used for subsequent analysis. These regressors were selected based on a systematic assessment comparing different confound models (Speer et al., 2024).

Since we were interested in changes in mental state that do not operate on a second-to-second scale, we applied temporal Gaussian smoothing of the signal to increase the signal-to-noise ratio to the preprocessed and cleaned data. Specifically, we applied a Gaussian smoothing kernel with a width of six TRs (9s) to the data. The cleaned data were segmented into individual prompts to examine the trial effects over time. To prevent confounding our analysis with the BOLD response to the onset and offset of the discussion prompt and task switching, we truncated each trial by removing the first and last seven TRs (10.5s) as recommended by previous research (Nastase et al., 2019).

### Measures

#### Agreement

Our primary behavioral outcome was participants’ agreement on the solution to each discussion problem. We measured participants’ quantitative allocation of money across five solutions before and after each conversation. This measure reflects how much they valued each option and served as the opinion profiles for each participant. We calculated the Mahalanobis distance between the profiles of the two dyad members before the conversation and after the conversation. We then multiplied the distance by −1 so that higher values equaled more agreement. We compared levels of agreement before and after the conversation to determine if discussions increased or decreased consensus and how this differed across conditions.

#### Exploration

In this study, we operationalized exploration in two ways. First, we looked at the distance between conversation partners over time, which allowed us to examine to what extent they *diverge* as the conversation progresses. Second, we investigated how much *novelty* the conversations contain. Novelty reflects the overall extent of territory explored, while divergence tracks changes in the relative distance between the two conversation partners. Exploratory dyads will move farther apart (rather than closer together) within the space and cover more diverse territory.

We examined exploration in two spaces:

**Mental State Space.** Using both neural and linguistic data, we assessed how pairs navigated a three-dimensional space of mental states—defined by social impact, rationality, and valence (the “3D mind model”; Tamir et al., 2016; Tamir & Thornton, 2018). A person’s location in this space reflects their internal state, and how two individuals move relative to each other reveals their interpersonal dynamics. Exploratory pairs tend to diverge within this space, moving farther apart and covering more varied mental states.

**Topic Space.** We also studied the exploration of conversation topics through linguistic analysis. Topic modeling revealed what dyads discussed over time. Exploratory behavior here was indicated by covering a wider range of topics, switching topics more often, and making more significant leaps between topics. Each of these measures reflects how broadly and dynamically the dyads explored conversational content.

#### Mental State Exploration

**Individual Locations.** To measure mental state exploration, both as divergence and novelty, we needed to first locate each participant’s mental state at each moment in the conversation.

For the neural data, we applied decoding models—originally trained on an independent dataset—to identify mental state locations from whole-brain activity patterns (Speer et al., 2024). In a previous study, Speer and colleagues (2024) trained machine learning models to decode mental state representations from whole-brain neural patterns recorded while participants completed tasks that elicited mental states varying along three key dimensions. Building on these models, we decoded each participant’s “location” in this mental state space during the conversation. Specifically, at every time point, we used the decoding models to analyze each conversation partner’s whole-brain activity, producing a time series of mental state coordinates in three-dimensional space. We then used these coordinates to calculate the distance between the two partners, as described below under **dyadic divergence** and **dyadic novelty**.

We analyzed participants’ mental state locations from their language as well, using a natural language processing (NLP) algorithm, the *affectR* package (https://github.com/markallenthornton/affectr), to classify words along the same three mental state dimensions. By extending an initial set of 166 rated mental state words (collected from 1,205 participants) to a two-million-word dictionary using fastText embeddings and a support vector regression model, we were able to generalize this three-dimensional analysis to virtually any word (see Supplementary Material 1 for more detail). This tool allowed us to identify which states people are experiencing based on the words they said and also enabled us to calculate how far apart two conversation partners are in that linguistic mental state space—shedding light on how closely their thoughts and feelings align. This enabled us to measure **dyadic divergence** and **dyadic novelty** in language.

**Dyadic Divergence.** For the neural data, we used the values for the three dimensions for each subject and time point to compute the Mahalanobis distance between the two conversation partners at each time point (TR). This gave us a measure of how distant the two partners are in mental state space relative to each other. Combining the relative distance measure with time, we were able to measure how close (or far) two conversation partners are to each as they enter the conversation and how the partners converge (or diverge) as they converse with each other. The Mahalanobis distance was chosen due to its insensitivity to the scale of the variables entered and because it removes redundant information from correlated variables. As a consequence, it has become the standard distance measure employed for multivariate fMRI analyses (Allefeld & Haynes, 2014; Bobadilla-Suarez et al., 2020; Nili et al., 2014).

For the linguistic data, we used the same approach with the only difference that we computed the distance in mental state space between each turn within a conversation. We exclusively used trials with more than three turns (two trials were removed), because to estimate the Mahalanobis distance, the number of samples must exceed the number of dimensions (*N* = 3: rationality, social impact & valence) to be able to calculate the covariance matrix needed. Participants signaled the end of their turn by pressing a button, facilitating the demarcation of turns. Trials with an excessive number of turns were excluded to ensure meaningful analysis with NLP tools. The 1.5 x interquartile range rule identified trials with over 21 turns as outliers, resulting in the removal of these trials (four trials, < 4%) from the NLP analysis. This resulted in the exclusion of one participant. Additionally, the number of words per turn was included as a nuisance predictor in regression models (detailed below). This was done to be able to account for the effect of length of turns on conversation outcomes.

**Dyadic Novelty.** To test how much dyads explore mental state space as they are engaged in a conversation, we adapted an approach from a previous study that investigated movement synchrony (Ravreby et al., 2022). This method tests how different (i.e., non-repetitive) each conversation turn is to the preceding turns, and accordingly, assesses a dyad’s mental state *novelty*. Novelty and divergence tap into different components of exploration: Novelty reflects how much new ground the conversation covers, whereas divergence reflects how far apart the two partners are in that space. It’s possible for two conversation partners to remain significantly distant from each other (high divergence) without exploring new areas (low novelty)—they may occupy separate positions in familiar territory without venturing into uncharted topics. As a consequence, we decided to measure both components of exploration.

For the neural data, we used the Kolmogoriv-Smirnov distance (K-S distance; Massey, 1951) to compute the distance between the empirical distribution of the Z-scored mental state locations of both partners of a given conversation turn to each of the previous conversation turns. So, for each conversation turn, the distribution of mental state locations of both conversation partners was compared to all the distributions of previous turns. All turns with less than three timepoints were removed from the analysis (6% of all turns). Each test involves comparing two turns at a time. The two-sample K-S test, which checks if two samples originate from the same distribution, is particularly appropriate for our situation because it is sensitive to differences in both the location and shape of the empirical cumulative distribution functions (CDFs). The distance between the empirical CDFs is small when the distributions are similar (it is zero when they are identical) and approaches one when they are very different. For each dyad, we compared the first and the second turns using the K-S distance test. Since there are no turns before the first one, we also assigned it as the novelty score of the first segment. Subsequently, we compared the third turn to each of the previous turns: we compared the third turn and the second turn, and the third turn and the first turn. The average statistic of the K-S distance tests of the two comparisons served as the dyadic novelty score of the third segment. This procedure was repeated for each dyad up to the last turn N. Thus, each turn was assigned a dyadic novelty score, which allowed us to track the novelty changes along the time domain. In order to have a single dyadic novelty score for each participant, we averaged the exploration scores of all the turns. This enabled us to test how much new ground a dyad covers in *neural* mental state space in absolute terms.

To measure novelty in linguistic mental state space, the same approach as above was used with the difference that we were only able to extract mental state locations of the person speaking. So, here we compared distributions of words between sentences instead of turns. Each sentence had an average of 21 words (SD = 14) with a minimum of three words, which made it suitable for our analysis. For each trial, we extracted the mental state location of each word the speaker said. We then compared the distribution of mental state locations of each new sentence to all the previous sentences as outlined above. This allowed us to test how much new ground a dyad covers in *linguistic* mental state space in absolute terms.

#### Topic Exploration

**Individual Locations.** The previous analyses extracted the conversation partners’ locations in mental state space but did not delve into the topics discussed. To tackle this question, we employed topic modeling to investigate the relationship between neural and linguistic mental dynamics and conversation content. Using BERTopic, we extracted topic representations for each sentence in every conversation across all dyads (Supplementary Material 2).

**Dyadic Divergence.** We calculated the distance in topic space from one turn to the next using the cosine distance. We adopted an approach used in a previous hyperscanning study focusing on affiliative conversations (Speer et al., 2024): the choice of cosine distance was motivated by the high-dimensional embedding space (>300 dimensions) in which topics are encoded, surpassing the number of samples (in this case, participants), rendering the computation of Mahalanobis distance unfeasible. Cosine distance was chosen over Euclidean distance because the Euclidean distance encounters problems in high-dimensional space (Aggarwal et al., 2001). In addition, the cosine similarity has been established as the standard distance measure for NLP. It is also the default metric in BERTopic because it does not take into account the magnitude of vectors, which is helpful when working with text data where word counts are influenced by the length of the documents, which often vary considerably in NLP analyses.

**Dyadic Novelty.** We used BERTopic to label each sentence that a conversation partner said with a topic. These topics were represented by numerical embeddings that are represented in a high-dimensional embedding space (>300 dimensions). This allowed us to generate a list of the topics discussed by a given dyad and their respective location in embedding space. We then computed dyadic novelty as follows: As a first step, we computed the pairwise cosine distance between each topic within a dyad. This told us how far each topic was from each of the other topics discussed by a given dyad. We then averaged these distances to obtain a single value representing topic novelty. This allowed us to gauge the extent of content space exploration by each dyad.

**Number of Topics.** Using the list of topics and their locations in high-dimensional embedding space for each dyad also allowed us to quantify the number of topics generated. To this end, we counted the number of unique topics each dyad discussed. This provided us with an additional measure of the scope of topic exploration.

**Topic Switches.** BERTopic also provided us with the sequence of topics discussed by each dyad. Each sequence of sentences within a dyad had a sequence of labels of the associated topics. This allowed us to examine the frequency of transitions between topics, including shifts back to previously discussed topics within a conversation segment. This provided a third measure of topic exploration.

### Analyses

#### Do Social Motives Influence Agreement?

We tested whether conversations increase agreement. We entered the dyads’ distance in opinion before and after the conversation into a multilevel (discussion problems nested in dyads) two-sample t-test to test whether dyads agreed more before or after the conversation. Next, we added the social motive as a predictor to the model to investigate whether there are differences in how much dyads align their opinions contingent on whether they were instructed to compromise or persuade.

#### Do Social Motives Influence Conversation Dynamics?

**Dyadic Divergence.** To test how the instruction to compromise vs. persuade shaped exploration in mental state and topic space, we fit regression models to our data. Given the nested structure of our data (time points within trials within participants), we conducted multilevel analyses to test the effect of social motive (instruction to *persuade* vs. *compromise*) and time (time points and trials) on exploration. Since each discussion problem constituted an independent conversation, we were particularly interested in time points (for neural data) and turns (for linguistic and topic data) within each prompt. Thus, in combination with social motive, time points within trials were considered the primary independent variable. The dependent variable for the neural and linguistic data was the continuous Mahalanobis distance with a Gaussian link representing distance in mental state space between the conversation partners. For the topic data, the dependent variable was the continuous cosine distance representing the distance in topic space with a Gaussian link. The social motive condition served as a dyad-level predictor, whereas time served as a trial-level (discussion problem) and a within-trial-level (time points for neural data and turns for topic and linguistic data) predictor. The models allowed for random intercepts within participants. The multilevel models were implemented in R (R Core Team, 2022) using the *nlme* package (Pinhero & Bates, 2023). This modeling approach allowed us to test how the instruction to compromise vs. persuade shapes exploration of content and mental state space in conversation over time and how the different measures relate to each other.

**Dyadic Novelty.** Given the nested structure of our data (trials within participants), we applied a multilevel two-sample t-test to test whether our measures of dyadic mental state and topic novelty differ between dyads instructed to compromise and persuade. This allowed us to test whether there are differences in absolute exploration between dyads instructed to compromise and persuade.

#### Do Conversation Dynamics Influence Agreement?

Next, we examined how exploration (divergence and novelty) in mental state and topic space predicted agreement.

**Dyadic Divergence.** To examine how agreement affects divergence, we fit three multilevel regression models to predict each of the three distance metrics, respectively: Mahalanobis distance in neural and linguistic mental state space and cosine distance in topic space. Each model includes agreement, trial number, and time points (within-dyads) as predictors and also controlled for initial difference in opinion (distance in opinion space before the conversation). This allowed us to test how eventual agreement influences how dyads traverse mental state and topic space relative to each other.

**Dyadic Novelty.** To test how novelty is associated with agreement, we fit three different multilevel regression models to predict agreement. Specifically, we fit separate models with neural, linguistic, and topic novelty, each also containing the initial distance in opinion (initial agreement) as nuisance regressors. In addition, we also fit two multilevel models to test for the effect of the number of topics generated and the number of topic switches on agreement, also controlling for initial agreement. This allowed us to test how absolute novelty in mental state and topic space predicts agreement.

#### Do Conversation Dynamics Mediate the Effect of Social Motives on Agreement?

In addition, we wanted to see how the effect of social motives (compromise vs. persuade) on agreement is mediated by exploration. To address this, we created a composite measure of exploration by calculating the mean over the neural, linguistic mental state, and topic novelty. This resulted in a composite exploration score for each trial for each dyad. We used a composite across the three measures because analyses revealed that all three modalities (neural, linguistic, and topic) tap into the same underlying construct (see Supplementary Material 3) and because the composite measure best predicts agreement (see Supplementary Material 4). That said, mediation analyses with each individual measure of exploration replicated this same pattern of results (Supplementary Material 5). Second, we used the novelty measure because it offers a more intuitively interpretable metric (i.e., absolute exploration) than divergence over time (see Discussion) and because it was found to be more predictive of agreement at the end of the conversation (see Supplementary Material 6). To examine the potential mediating effect of exploration on the relationship between conversation goal and agreement, we conducted a mediation analysis using the *mediate* function from the *mediation* package in R (Tingley et al., 2014).

We used nonparametric bootstrapping with 5,000 simulations to estimate confidence intervals for the indirect effect. The average causal mediation effect (ACME) and the average direct effect (ADE) were calculated, along with the total effect. Statistical significance of the mediation was assessed using 95% confidence intervals. All analyses were performed in R, version 4.2.1.

## Supplementary Material

### Supplementary Material 1: AffectR Algorithm

We implemented the affectR package to decode how dyads traverse linguistic mental state space based on the words they chose to say in the conversation. *affectR* allows us to measure the sentiment of words along the three mental state dimensions (rationality, social impact, and valence). The *affectR* package was created using ratings of mental state words on 16 psychological dimensions (Tamir et al., 2016). The ratings were collected from 1205 participants on Amazon Mechanical Turk, with aggregate values freely available online (https://osf.io/3qn47/). Principal component analysis (PCA) revealed that these 16 dimensions could be reduced to three mental state dimensions, which proved able to explain patterns of brain activity associated with thinking about others’ mental states (Tamir et al., 2016). To expand the volume of the dictionary from 166 words to two million words, the fastText library (https://fasttext.cc/) was employed. A support vector regression (SVR) model with a radial basis function was used to predict the ratings of 166 mental state words along three principal component dimensions. This approach was feasible because these 166 words are part of the fastText corpus. The SVR model was then used to estimate 3D dimension scores for all 2 million words in the fastText corpus, resulting in the creation of a comprehensive weighted dictionary. This approach has been implemented in a previous neuroimaging study (Thornton & Tamir, 2020) to show that the 3D mind model does not capture how mental states are represented only in the brain but also in language. Further, in more recent studies, the same method was applied to study whether the 3D mind model characterizes how people understand mental states across modern and historical cultures (Thornton et al., 2022) and to explore what mental state trajectories lead to stronger social connection in affiliative conversations (Speer et al., 2024).

### Supplementary Material 2: Topic Modeling Using BERTopic

We applied topic modeling to explore how the neural and linguistic dynamics relate to the content of the conversations. Topic modeling is gaining popularity in psychology and social neuroscience. It has previously been successfully used in the domain of marketing on social media conversations to drive brand strategy (Resnik et al., 2015) and has been implemented in clinical psychology to differentiate between the language of depressed and non-depressed individuals (Putnick & Bornstein, 2016) and to detect self-injurious thoughts and behavior on online platforms (Franz et al., 2020). We employed BERTopic (https://maartengr.github.io/BERTopic/index.html), which is a topic modeling technique that leverages hugging-face (https://huggingface.co/) transformers and class-based term frequency-inverse document frequency (cTF-DF) to create dense clusters providing easily interpretable topics while keeping important words in the topic descriptions. The benefit of BERTopic, as opposed to more traditional algorithms, is that it selects the number of topics automatically and does not require the specification of number of topics beforehand, which enables a purely data-driven exploration of topics. In addition, BERTopic does not only rely on word-level but also uses sentence-level embeddings, which has the advantage of considering the semantic relationship between words in a corpus, leading to more informative topics. Another advantage is that due to the nature of the semantic embeddings, textual preprocessing (stemming, lemmatization, stopword removal, etc.) is not needed.

### Supplementary Material 3: Correlation Between Exploration Across Modalities

To test whether there is internal consistency between the neural, linguistic, and topic exploration measures (novelty and divergence), we computed the Pearson correlation between all six measures. To obtain measures for neural, linguistic, and topic divergence, we extracted the random slopes for each participant from the three multilevel models depicted in Figure 3. That is, the multilevel model fit on the neural data, the linguistic data, and the topic modeling data. The correlation analysis revealed that the six measures of exploration are overall positively correlated, which suggests that our measures of exploration are internally consistent (see Supplementary Figure 1). All three measures of novelty were also strongly and significantly positively correlated. Further, the divergence measures showed a positive correlation. The observation that the neural and linguistic divergence are correlated was expected because these two measures exist in the same mental state space, and the observation that linguistic and topic divergence are correlated was anticipated because they are both based on linguistic data. As a result, the fact that the correlation between topic and neural divergence was not significant was expected to reveal the weakest correlation because the conceptual and methodological distance was the largest. We also observed significant correlations between topic novelty and linguistic and neural divergence, which was expected because topic novelty was also based on distances of individual data points, whereas neural and linguistic novelty were based on comparisons between distributions. We also observed a marginally significant correlation between linguistic divergence and neural and linguistic novelty. Together, these results suggest that our measures of exploration are internally consistent.

**Supplementary Figure 1.**
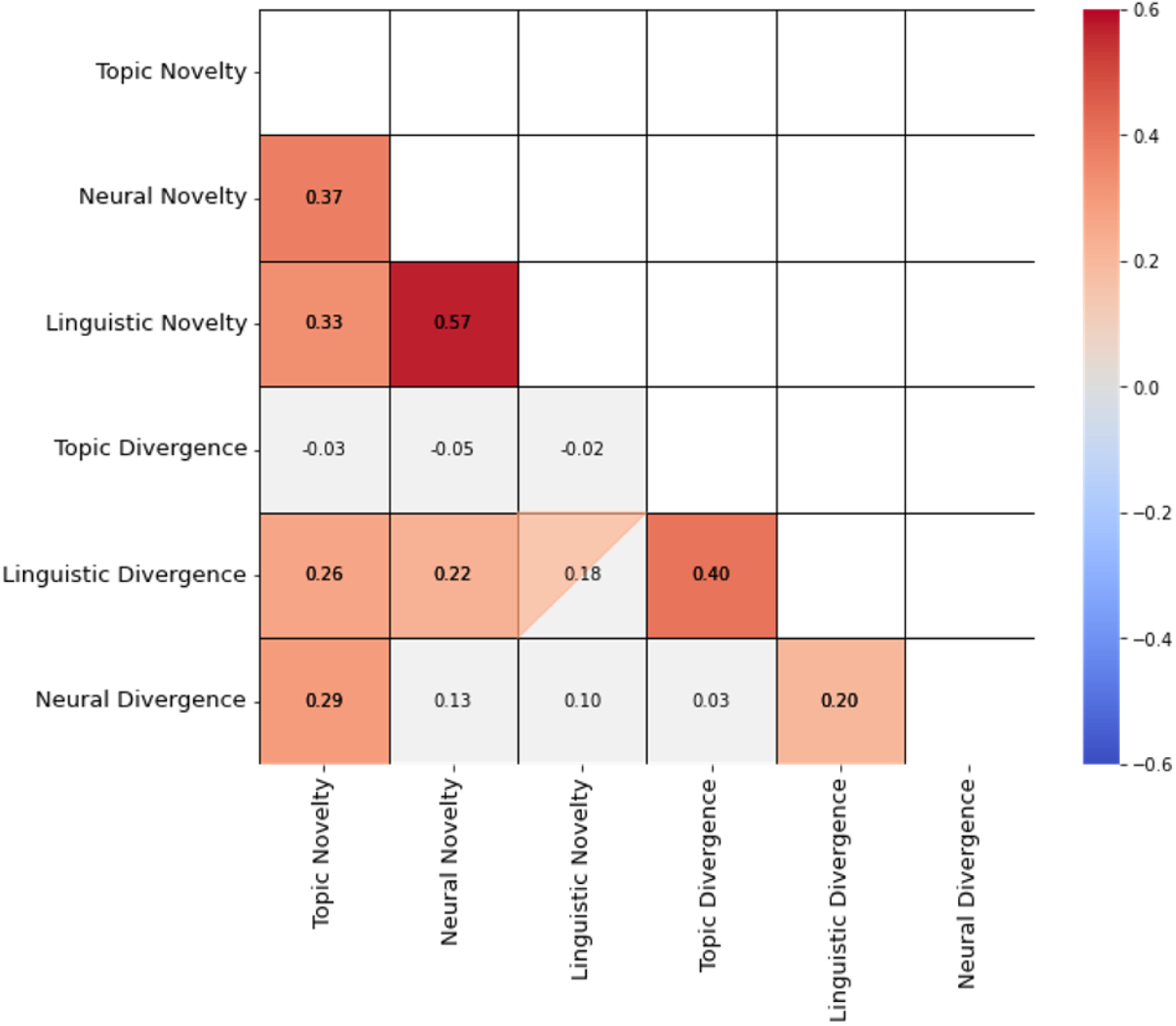
Correlation between neural, linguistic and topic novelty and divergence. Colored squared are significant at *p* < 0.05. Squares that are filled with a triangle are marginally significant (*p* < 0.10). Grey squares are not significant.

### Supplementary Material 4: Individual vs. Dyadic Exploration aAcross Modalities

We examined whether exploration is mostly driven by one of the partners being exploratory or whether there are synergistic effects of both partners exploring. To this end, we performed a model comparison where we fit a model with only individual exploration and a model with only dyadic exploration predicting agreement (while also controlling for the number of turns and the initial distance in opinions). Individual neural and linguistic exploration were computed the same way as the dyadic neural novelty with the only difference that here we only focused on the turns of one of the participants at a time. For individual topic exploration, the mean distance between topics introduced by one participant at a time was computed. To compare the models, we used the Bayesian Information Criterion (BIC; lower values represent higher fit). The model comparison revealed that for all measures of exploration, except for topic exploration, dyadic exploration was a better predictor than the two individual partners’ exploration, suggesting that the combined dyadic exploration is greater than the sum of its parts and highlighting the synergistic effects of dyadic exploration.

This analysis also revealed that the dyadic neural and linguistic exploration models predict agreement equally well, whereas the topic exploration measure does a poorer job at predicting agreement. Further, these models explain different portions of the variance in agreement across dyads, as the composite measure of all three of these measures does substantially better in predicting agreement than each individual modality.

**Supplementary Figure 2.**
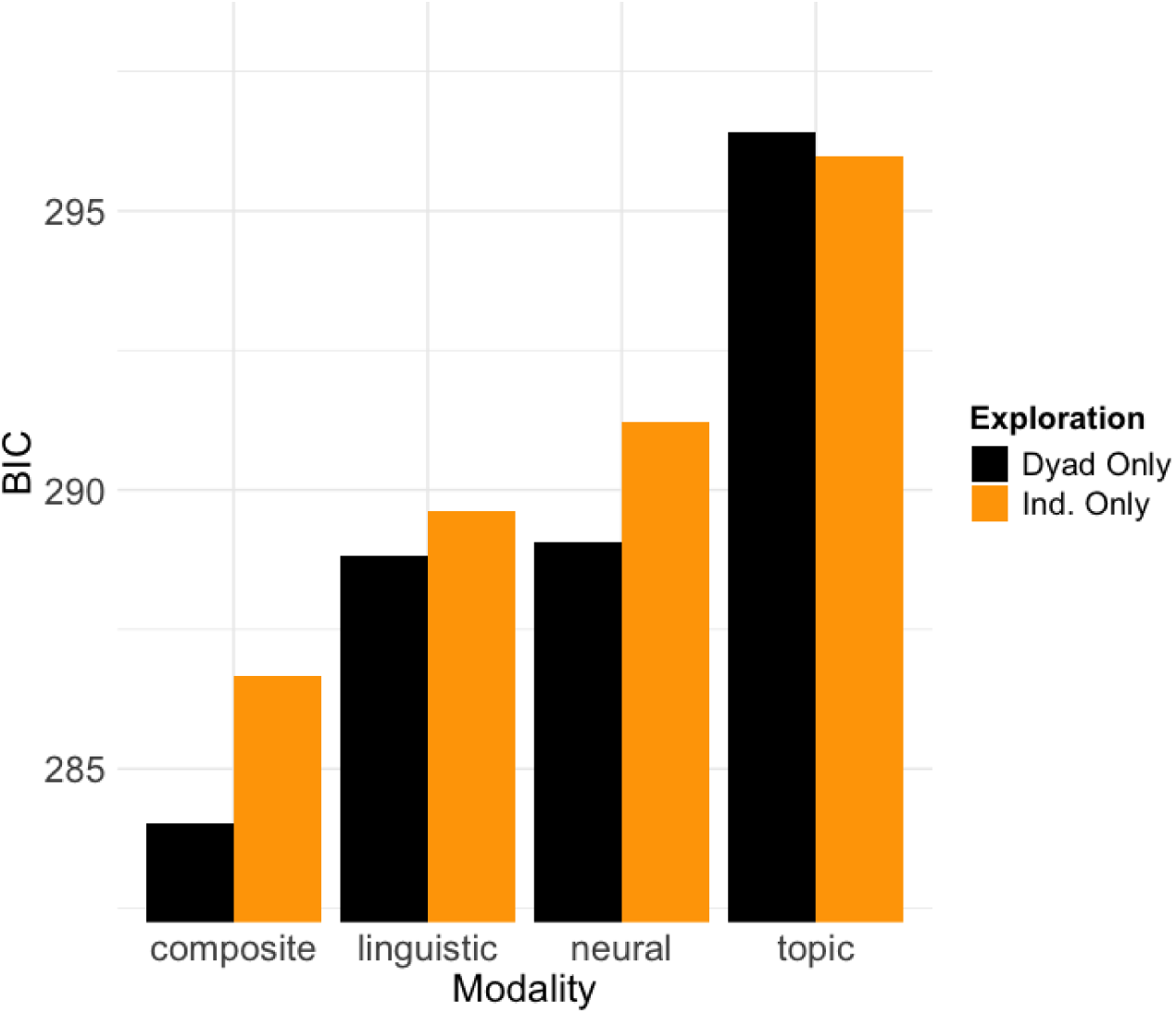
Dyadic exploration is a better predictor than individual exploration. This graph shows the model fit (BIC; lower is better) for a model with both participants’ individual exploration or their dyadic exploration as a predictor of agreement across the 3 modalities and their composite.

### Supplementary Material 5: Mediation Analyses Robustness Check

To test the robustness of our mediation analysis, we also conducted a mediation analysis with each individual measure of exploration in terms of novelty as the mediation. The analysis revealed a similar pattern as the findings in Supplementary Material 3: the mediation analysis revealed significant and marginally significant mediation effects for the linguistic and neural mental state exploration measures, respectively. No significant mediation was found for the topic exploration measure, where only the effect of social motive on exploration was significant (see Supplementary Figure 2). In addition, these analyses showed that these measures appear to capture different portions of variance in dyads’ exploration behavior, as the composite measure performs best in predicting agreement and is the only one that fully mediates the effect of social motive on agreement. No significant mediation effects were found for the divergence measure of exploration.

**Supplementary Figure 3.**
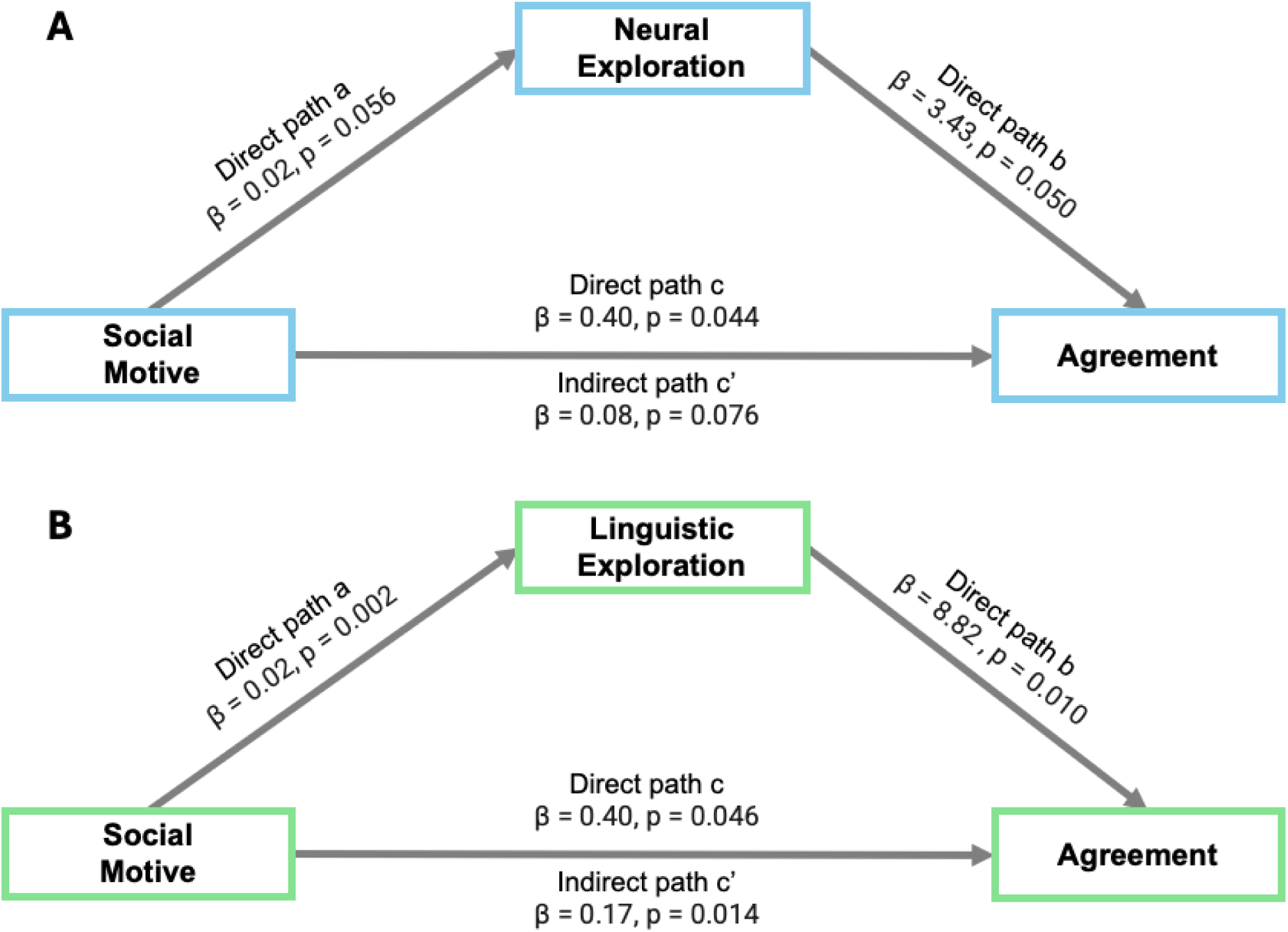
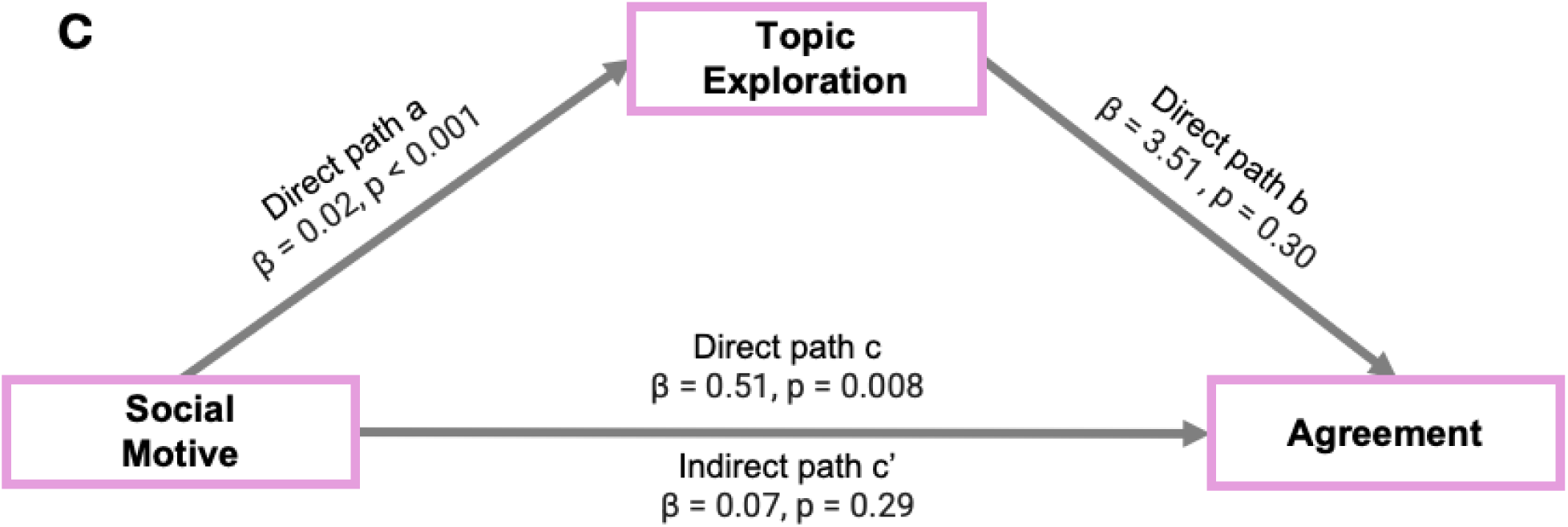
Exploration mediates the positive effect of compromise on agreement. Mediation analyses for the neural A), linguistic B) and topic C) novelty measures.

### Supplementary Material 6: Comparison Between Divergence and Novelty in Predicting Eventual Agreement

We also tested whether divergence or novelty is more important in predicting eventual agreement. We used the slopes of a multilevel model predicting distance between partners (in neural, linguistic, or topic space) from time as a measure of divergence. For each modality (neural, linguistic and topic), we then entered the divergence and novelty measures into a multilevel model predicting eventual agreement while controlling for initial agreement. For both neural (β_novelty_ = 4.05, 95% CI [1.81, 6.27], SE = 1.11, p < 0.001) and linguistic (β_novelty_ = 10.79, 95% CI [4.92, 16.66], SE = 2.92, p < 0.001) data, only the novelty measure significantly predicted eventual agreement. For the topic data (β_novelty_ = 5.97, 95% CI [-0.75, 12.69], SE = 3.34, p = 0.08) there was a marginally significant effect of novelty. In none of the models did divergence reach significance as a predictor of eventual agreement. In addition, divergence was not found to be a significant mediator of social motive on agreement. This suggests that novelty is more important in predicting agreement at the end of the conversation as opposed to divergence.

### Supplementary Material 7: Dyads Who Agree More Before the Conversation Diverge More

To test whether our measures of exploration relate to the initial agreement, that is, the inverse distance in opinion at the start of the conversation, we computed the correlation between all six of our exploration measures (divergence & novelty for neural, linguistic, and topic data). We found a significant correlation between neural divergence and initial agreement (*r* = 0.22, *p* = 0.027) and a marginally significant correlation between topic divergence and initial agreement (*r* = 0.19, *p* = 0.062). No significant correlation was found between linguistic divergence and initial agreement. When combining the neural and topic divergence measure using their first principal component, a strong positive correlation was found (*r* = 0.30, *p* = 0.003). No significant correlations were found between the novelty measures and initial alignment. This suggests that dyads who enter the conversations with more aligned opinions are more inclined to diverge in mental state and topic space.

**Supplementary Figure 4.**
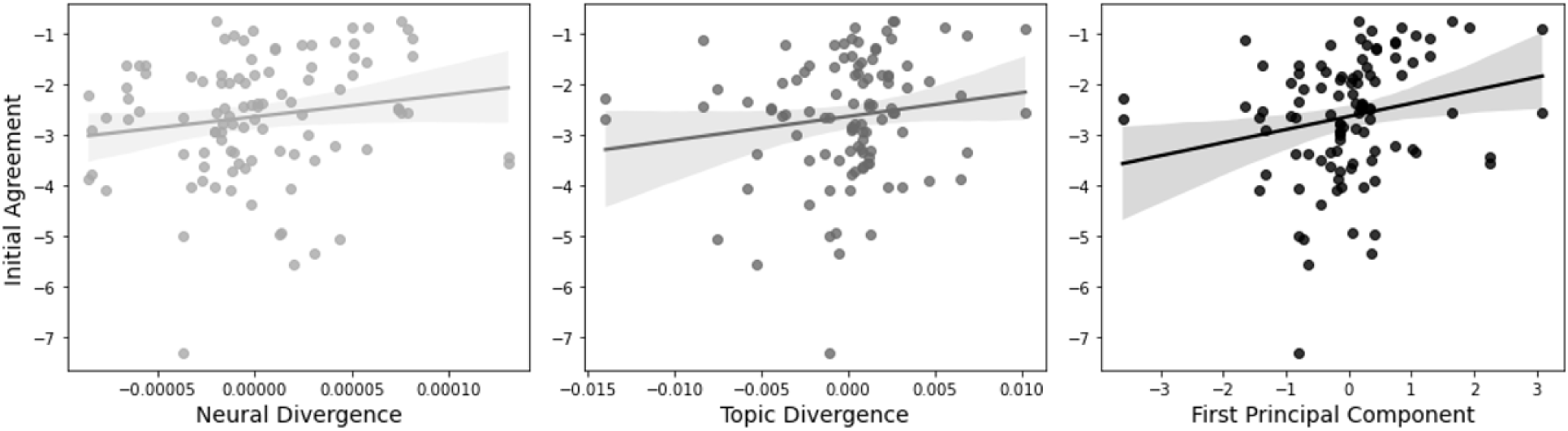
Neural and topic divergence is positively correlated with agreement when entering the conversation.

## References

Abney, D. H., Paxton, A., Dale, R., & Kello, C. T. (2015). Movement dynamics reflect a functional role for weak coupling and role structure in dyadic problem solving. Cognitive Processing, 16(4), 325–332. 10.1007/s10339-015-0648-2

Aggarwal, C. C., Hinneburg, A., & Keim, D. A. (2001). On the Surprising Behavior of Distance Metrics in High Dimensional Space. In J. Van den Bussche & V. Vianu (Eds.), Database Theory—ICDT 2001 (Vol. 1973, pp. 420–434). Springer Berlin Heidelberg. 10.1007/3-540-44503-X_27

Allefeld, C., & Haynes, J.-D. (2014). Searchlight-based multi-voxel pattern analysis of fMRI by cross-validated MANOVA. NeuroImage, 89, 345–357. 10.1016/j.neuroimage.2013.11.043

Aron, A., Melinat, E., Aron, E., Vallone, R., & Bator, R. (1997). The experimental generation of interpersonal closeness: A procedure and some preliminary findings. Personality and Social Psychology Bulletin, 23(4), 363–377.

Babiloni, F., & Astolfi, L. (2014). Social neuroscience and hyperscanning techniques: Past, present and future. Neuroscience & Biobehavioral Reviews, 44, 76–93. 10.1016/j.neubiorev.2012.07.006

Balconi, M. (2010). The Sense of Agency in Psychology and Neuropsychology. In B. Michela (Ed.), Neuropsychology of the Sense of Agency: From Consciousness to Action (pp. 3–22). Springer Milan. 10.1007/978-88-470-1587-6_1

Beersma, B., & De Dreu, C. K. W. (2002). Integrative and Distributive Negotiation in Small Groups: Effects of Task Structure, Decision Rule, and Social Motive. Organizational Behavior and Human Decision Processes, 87(2), 227–252. 10.1006/obhd.2001.2964

Behrens, F., Snijdewint, J. A., Moulder, R. G., Prochazkova, E., Sjak-Shie, E. E., Boker, S. M., & Kret, M. E. (2020). Physiological synchrony is associated with cooperative success in real-life interactions. Scientific Reports, 10(1), 19609. 10.1038/s41598-020-76539-8

Bittner, J. V., Bruena, M., & Rietzschel, E. F. (2016). Cooperation goals, regulatory focus, and their combined effects on creativity. Thinking Skills and Creativity, 19, 260–268. 10.1016/j.tsc.2015.12.002

Bobadilla-Suarez, S., Ahlheim, C., Mehrotra, A., Panos, A., & Love, B. C. (2020). Measures of Neural Similarity. Computational Brain & Behavior, 3(4), 369–383. 10.1007/s42113-019-00068-5

Brett, J. M. (2000). Culture and Negotiation. International Journal of Psychology, 35(2), 97–104. 10.1080/002075900399385

Brett, J. M., Shapiro, D. L., & Lytle, A. L. (1998). BREAKING THE BONDS OF RECIPROCITY IN NEGOTIATIONS. Academy of Management Journal, 41(4), 410–424. 10.2307/257081

Brett, J., & Thompson, L. (2016). Negotiation. Organizational Behavior and Human Decision Processes, 136, 68–79. 10.1016/j.obhdp.2016.06.003

Brooks, A. W., Gino, F., & Schweitzer, M. E. (2015). Smart People Ask for (My) Advice: Seeking Advice Boosts Perceptions of Competence. Management Science, 61(6), 1421–1435. 10.1287/mnsc.2014.2054

Brynjarsdottir, H., Håkansson, M., Pierce, J., Baumer, E., DiSalvo, C., & Sengers, P. (2012). Sustainably unpersuaded: How persuasion narrows our vision of sustainability. Proceedings of the SIGCHI Conference on Human Factors in Computing Systems, 947–956. 10.1145/2207676.2208539

Burns, S. M., Tsoi, L., Falk, E. B., Speer, S. P. H., & Tamir, D. I. (2024). Interdependent minds: Quantifying the dynamics of successful social interaction. PsyArXiv.

Carmeli, A., Dutton, J. E., & Hardin, A. E. (2015). Respect as an engine for new ideas: Linking respectful engagement, relational information processing and creativity among employees and teams. Human Relations, 68(6), 1021–1047. 10.1177/0018726714550256

Cooney, G., Gilbert, D. T., & Wilson, T. D. (2017). The Novelty Penalty: Why Do People Like Talking About New Experiences but Hearing About Old Ones? Psychological Science, 28(3), 380–394. 10.1177/0956797616685870

De Dreu, C. K. W., Evers, A., Beersma, B., Kluwer, E. S., & Nauta, A. (2001). A theory-based measure of conflict management strategies in the workplace. Journal of Organizational Behavior, 22(6), 645–668. 10.1002/job.107

De Dreu, C. K. W., Weingart, L. R., & Kwon, S. (2000). Influence of social motives on integrative negotiation: A meta-analytic review and test of two theories. Journal of Personality and Social Psychology, 78(5), 889–905. 10.1037/0022-3514.78.5.889

Decety, J., Jackson, P. L., Sommerville, J. A., Chaminade, T., & Meltzoff, A. N. (2004). The neural bases of cooperation and competition: An fMRI investigation. NeuroImage, 23(2), 744–751. 10.1016/j.neuroimage.2004.05.025

Deutsch, M. (1973). The resolution of conflict. Yale University Press.

Dideriksen, C., Christiansen, M. H., Tylén, K., Dingemanse, M., & Fusaroli, R. (2023). Quantifying the Interplay of Conversational Devices in Building Mutual Understanding.

Doré, B. P., & Morris, R. R. (2018). Linguistic Synchrony Predicts the Immediate and Lasting Impact of Text-Based Emotional Support. Psychological Science, 29(10), 1716–1723. 10.1177/0956797618779971

Druckman, D. (1986). Stages, Turning Points, and Crises: Negotiating Military Base Rights, Spain and the United States. Journal of Conflict Resolution, 30(2), 327–360. 10.1177/0022002786030002006

Fisher, M., Knobe, J., Strickland, B., & Keil, F. C. (2017). The Influence of Social Interaction on Intuitions of Objectivity and Subjectivity. Cognitive Science, 41(4), 1119–1134. 10.1111/cogs.12380

Fisher, R., Ury, W., & Patton, B. (2011). Getting to yes: Negotiating agreement without giving in. Penguin.

Franz, P. J., Nook, E. C., Mair, P., & Nock, M. K. (2020). Using Topic Modeling to Detect and Describe Self-Injurious and Related Content on a Large-Scale Digital Platform. Suicide and Life-Threatening Behavior, 50(1), 5–18. 10.1111/sltb.12569

Gelfo, F. (2019). Does Experience Enhance Cognitive Flexibility? An Overview of the Evidence Provided by the Environmental Enrichment Studies. Frontiers in Behavioral Neuroscience, 13, 150. 10.3389/fnbeh.2019.00150

Gilson, L. L., & Shalley, C. E. (2004). A Little Creativity Goes a Long Way: An Examination of Teams’ Engagement in Creative Processes. Journal of Management, 30(4), 453–470. 10.1016/j.jm.2003.07.001

Goldstone, R. L., Andrade-Lotero, E. J., Hawkins, R. D., & Roberts, M. E. (2024). The Emergence of Specialized Roles Within Groups. Topics in Cognitive Science, 16(2), 257–281. 10.1111/tops.12644

Gouldner, A. W. (1960). The Norm of Reciprocity: A Preliminary Statement. American Sociological Review, 25(2), 161. 10.2307/2092623

Hargadon, A. B., & Bechky, B. A. (2006). When Collections of Creatives Become Creative Collectives: A Field Study of Problem Solving at Work. Organization Science, 17(4), 484–500. 10.1287/orsc.1060.0200

Hoehl, S., Fairhurst, M., & Schirmer, A. (2021). Interactional synchrony: Signals, mechanisms and benefits. Social Cognitive and Affective Neuroscience, 16(1–2), 5–18. 10.1093/scan/nsaa024

Hommel, B. (2015). Between Persistence and Flexibility. In Advances in Motivation Science (Vol. 2, pp. 33–67). Elsevier. 10.1016/bs.adms.2015.04.003

Hon, A. H. Y., Bloom, M., & Crant, J. M. (2014). Overcoming Resistance to Change and Enhancing Creative Performance. Journal of Management, 40(3), 919–941. 10.1177/0149206311415418

Ireland, M. E., Slatcher, R. B., Eastwick, P. W., Scissors, L. E., Finkel, E. J., & Pennebaker, J. W. (2011). Language Style Matching Predicts Relationship Initiation and Stability. Psychological Science, 22(1), 39–44. 10.1177/0956797610392928

Jacob, C., Guéguen, N., Martin, A., & Boulbry, G. (2011). Retail salespeople’s mimicry of customers: Effects on consumer behavior. Journal of Retailing and Consumer Services, 18(5), 381–388. 10.1016/j.jretconser.2010.11.006

Jain, A., Department of Computer Engineering, SVKMs NMIMS MPSTME Shirpur, Maharashtra, India, Kulkarni, G., Department of Computer Engineering, SVKMs NMIMS MPSTME Shirpur, Maharashtra, India, Shah, V., & Department of Computer Engineering, SVKMs NMIMS MPSTME Shirpur, Maharashtra, India. (2018). Natural Language Processing. International Journal of Computer Sciences and Engineering, 6(1), 161–167. 10.26438/ijcse/v6i1.161167

Jones, S. M. (2011). Supportive Listening. International Journal of Listening, 25(1–2), 85–103. 10.1080/10904018.2011.536475

Kelsen, B. A., Sumich, A., Kasabov, N., Liang, S. H. Y., & Wang, G. Y. (2022). What has social neuroscience learned from hyperscanning studies of spoken communication? A systematic review. Neuroscience & Biobehavioral Reviews, 132, 1249–1262. 10.1016/j.neubiorev.2020.09.008

King-Casas, B., Tomlin, D., Anen, C., Camerer, C. F., Quartz, S. R., & Montague, P. R. (2005). Getting to Know You: Reputation and Trust in a Two-Person Economic Exchange. Science, 308(5718), 78–83. 10.1126/science.1108062

Knowles, E. S., & Linn, J. A. (Eds.). (2004). Resistance and persuasion. Lawrence Erlbaum Associates.

Koban, L., Ramamoorthy, A., & Konvalinka, I. (2019). Why do we fall into sync with others? Interpersonal synchronization and the brain’s optimization principle. Social Neuroscience, 14(1), 1–9. 10.1080/17470919.2017.1400463

Kong, D. T., Dirks, K. T., & Ferrin, D. L. (2014). Interpersonal Trust within Negotiations: Meta-Analytic Evidence, Critical Contingencies, and Directions for Future Research. Academy of Management Journal, 57(5), 1235–1255. 10.5465/amj.2012.0461

Krueger, F., McCabe, K., Moll, J., Kriegeskorte, N., Zahn, R., Strenziok, M., Heinecke, A., & Grafman, J. (2007). Neural correlates of trust. Proceedings of the National Academy of Sciences, 104(50), 20084–20089. 10.1073/pnas.0710103104

Laureiro-Martínez, D., & Brusoni, S. (2018). Cognitive flexibility and adaptive decision-making: Evidence from a laboratory study of expert decision makers. Strategic Management Journal, 39(4), 1031–1058. 10.1002/smj.2774

Lee, D. S., & Fujita, K. (2023). From whom do people seek what type of support? A regulatory scope perspective. Journal of Personality and Social Psychology, 124(4), 796–811. 10.1037/pspi0000405

Lewicki, R., Saunders, D., & Barry, B. (2024). Negotiation (9th ed.). McGraw-Hill.

Liberman, N., & Trope, Y. (2014). Traversing psychological distance. Trends in Cognitive Sciences, 18(7), 364–369. 10.1016/j.tics.2014.03.001

Liu, T., Saito, G., Lin, C., & Saito, H. (2017). Inter-brain network underlying turn-based cooperation and competition: A hyperscanning study using near-infrared spectroscopy. Scientific Reports, 7(1), 8684. 10.1038/s41598-017-09226-w

Liu, T., Saito, H., & Oi, M. (2015). Role of the right inferior frontal gyrus in turn-based cooperation and competition: A near-infrared spectroscopy study. Brain and Cognition, 99, 17–23. 10.1016/j.bandc.2015.07.001

Lu, K., Xue, H., Nozawa, T., & Hao, N. (2019). Cooperation Makes a Group be More Creative. Cerebral Cortex, 29(8), 3457–3470. 10.1093/cercor/bhy215

Ludwig, S., de Ruyter, K., Friedman, M., Brüggen, E. C., Wetzels, M., & Pfann, G. (2013). More than Words: The Influence of Affective Content and Linguistic Style Matches in Online Reviews on Conversion Rates. Journal of Marketing, 77(1), 87–103. 10.1509/jm.11.0560

Manson, J. H., Bryant, G. A., Gervais, M. M., & Kline, M. A. (2013). Convergence of speech rate in conversation predicts cooperation. Evolution and Human Behavior, 34(6), 419–426. 10.1016/j.evolhumbehav.2013.08.001

Massey, F. J. (1951). The Kolmogorov-Smirnov Test for Goodness of Fit. Journal of the American Statistical Association, 46(253), 68–78. 10.1080/01621459.1951.10500769

Mehl, M. R., Vazire, S., Holleran, S. E., & Clark, C. S. (2010). Eavesdropping on Happiness: Well-Being Is Related to Having Less Small Talk and More Substantive Conversations. Psychological Science, 21(4), 539–541. 10.1177/0956797610362675

Meina Liu, & Wilson, S. R. (2011). The Effects of Interaction Goals on Negotiation Tactics and Outcomes: A Dyad-Level Analysis Across Two Cultures. Communication Research, 38(2), 248–277. 10.1177/0093650210362680

Messick, D. M., & McClintock, C. G. (1968). Motivational bases of choice in experimental games. Journal of Experimental Social Psychology, 4(1), 1–25. 10.1016/0022-1031(68)90046-2

Nastase, S. A., Gazzola, V., Hasson, U., & Keysers, C. (2019). Measuring shared responses across subjects using intersubject correlation. Social Cognitive and Affective Neuroscience, nsz037. 10.1093/scan/nsz037

Nguyen, T., Kungl, M. T., Hoehl, S., White, L. O., & Vrtička, P. (2024). Visualizing the invisible tie: Linking parent–child neural synchrony to parents’ and children’s attachment representations. Developmental Science, e13504. 10.1111/desc.13504

Nili, H., Wingfield, C., Walther, A., Su, L., Marslen-Wilson, W., & Kriegeskorte, N. (2014). A Toolbox for Representational Similarity Analysis. PLoS Computational Biology, 10(4), e1003553. 10.1371/journal.pcbi.1003553

Olekalns, M., & Smith, P. L. (1999). Social Value Orientations and Strategy Choices in Competitive Negotiations. Personality and Social Psychology Bulletin, 25(6), 657–668. 10.1177/0146167299025006002

Olekalns, M., & Smith, P. L. (2003a). SOCIAL MOTIVES IN NEGOTIATION: THE RELATIONSHIPS BETWEEN DYAD COMPOSITION, NEGOTIATION PROCESSES AND OUTCOMES. International Journal of Conflict Management, 14(3/4), 233–254. 10.1108/eb022900

Olekalns, M., & Smith, P. L. (2003b). Testing the relationships among negotiators’ motivational orientations, strategy choices, and outcomes. Journal of Experimental Social Psychology, 39(2), 101–117. 10.1016/S0022-1031(02)00520-6

Perc, M., & Szolnoki, A. (2008). Social diversity and promotion of cooperation in the spatial prisoner’s dilemma game. Physical Review E, 77(1), 011904. 10.1103/PhysRevE.77.011904

Petty, R. E., & Cacioppo, J. T. (1986). Communication and persuasion: Central and peripheral routes to attitude change. Springer Science & Business Media.

Pinhero, J., & Bates, D. (2023). nlme: Linear and Nonlinear Mixed Effects Models. R Core Team. https://CRAN.R-project.org/package=nlme.

Powers, J. P., & LaBar, K. S. (2019). Regulating emotion through distancing: A taxonomy, neurocognitive model, and supporting meta-analysis. Neuroscience & Biobehavioral Reviews, 96, 155–173. 10.1016/j.neubiorev.2018.04.023

Pruitt, D., & Rubin, J. E. (1986). Social conflict: Escalation, stalemate and settlement. New York, NY: Random House.

Putman, W. B., & Street, R. L. (1984). The conception and perception of noncontent speech performance: Implications for speech-accommodation theory. International Journal of the Sociology of Language, 1984(46). 10.1515/ijsl.1984.46.97

Putnam, L. L., & Jones, T. S. (1982). Reciprocity in negotiations: An analysis of bargaining interaction. Communication Monographs, 49(3), 171–191. 10.1080/03637758209376080

Putnick, D. L., & Bornstein, M. H. (2016). Measurement invariance conventions and reporting: The state of the art and future directions for psychological research. Developmental Review, 41, 71–90. 10.1016/j.dr.2016.06.004

R Core Team. (2022). R: A language and environment for statistical computing. R Foundation for Statistical Computing. https://www.R-project.org/.

Ramseyer, F., & Tschacher, W. (2011). Nonverbal synchrony in psychotherapy: Coordinated body movement reflects relationship quality and outcome. Journal of Consulting and Clinical Psychology, 79(3), 284–295. 10.1037/a0023419

Ravreby, I., Shilat, Y., & Yeshurun, Y. (2022). Liking as a balance between synchronization, complexity and novelty. Scientific Reports, 12(1), 3181. 10.1038/s41598-022-06610-z

Reece, A. G., Cooney, G., Bull, P., Chung, C., Dawson, B., Fitzpatrick, C., Glazer, T., Knox, D., Liebscher, A., & Marin, S. (2022). *Advancing an Interdisciplinary Science of Conversation: Insights from a Large Multimodal Corpus of Human Speech* [Preprint]. PsyArXiv. 10.31234/osf.io/ts43f

Resnik, P., Armstrong, W., Claudino, L., Nguyen, T., Nguyen, V.-A., & Boyd-Graber, J. (2015). Beyond LDA: Exploring Supervised Topic Modeling for Depression-Related Language in Twitter. Proceedings of the 2nd Workshop on Computational Linguistics and Clinical Psychology: From Linguistic Signal to Clinical Reality, 99–107. 10.3115/v1/W15-1212

Sassenberg, K., & Winter, K. (2024). Intraindividual Conflicts Reduce the Polarization of Attitudes. Current Directions in Psychological Science.

Sened, H., Speer, S. P. H., Cooney, G., Reece, A. G., & Tamir, D. I. (2024). The Temporal Arc of Successful Conversations. Society for Personality and Social Psychology 2024 Conference, San Diego, CA.

Shaw, D. J., Czekóová, K., Staněk, R., Mareček, R., Urbánek, T., Špalek, J., Kopečková, L., Řezáč, J., & Brázdil, M. (2018). A dual-fMRI investigation of the iterated Ultimatum Game reveals that reciprocal behaviour is associated with neural alignment. Scientific Reports, 8(1), 10896. 10.1038/s41598-018-29233-9

Shirado, H., & Christakis, N. A. (2017). Locally noisy autonomous agents improve global human coordination in network experiments. Nature, 545(7654), 370–374. 10.1038/nature22332

Sievers, B., & Thornton, M. A. (2024). Deep social neuroscience: The promise and peril of using artificial neural networks to study the social brain. Social Cognitive and Affective Neuroscience, 19(1), nsae014. 10.1093/scan/nsae014

Slepian, M. L., & Moulton-Tetlock, E. (2019). Confiding Secrets and Well-Being. Social Psychological and Personality Science, 10(4), 472–484. 10.1177/1948550618765069

Speer, S. P. H., Mwilambwe-Tshilobo, L., Tsoi, L., Burns, S. M., Falk, E. B., & Tamir, D. I. (2024). Hyperscanning shows friends explore and strangers converge in conversation. Nature Communications, 15(1), 7781. 10.1038/s41467-024-51990-7

Street, R. L., Brady, R. M., & Putman, W. B. (1983). The Influence of Speech Rate Stereotypes and Rate Similarity or Listeners’ Evaluations of Speakers. Journal of Language and Social Psychology, 2(1), 37–56. 10.1177/0261927X8300200103

Su, Q., Li, A., Zhou, L., & Wang, L. (2016). Interactive diversity promotes the evolution of cooperation in structured populations. New Journal of Physics, 18(10), 103007. 10.1088/1367-2630/18/10/103007

Tamir, D. I., & Thornton, M. A. (2018). Modeling the Predictive Social Mind. Trends in Cognitive Sciences, 22(3), 201–212. 10.1016/j.tics.2017.12.005

Tamir, D. I., Thornton, M. A., Contreras, J. M., & Mitchell, J. P. (2016). Neural evidence that three dimensions organize mental state representation: Rationality, social impact, and valence. Proceedings of the National Academy of Sciences, 113(1), 194–199. 10.1073/pnas.1511905112

Thompson, L., & DeHarpport, T. (1994). Social Judgment, Feedback, and Interpersonal Learning in Negotiation. Organizational Behavior and Human Decision Processes, 58(3), 327–345. 10.1006/obhd.1994.1040

Thornton, M. A., & Tamir, D. I. (2020). People represent mental states in terms of rationality, social impact, and valence: Validating the 3d Mind Model. Cortex, 125, 44–59. 10.1016/j.cortex.2019.12.012

Thornton, M. A., Wolf, S., Reilly, B. J., Slingerland, E. G., & Tamir, D. I. (2022). The 3d Mind Model Characterizes How People Understand Mental States Across Modern and Historical Cultures. Affective Science, 3(1), 93–104. 10.1007/s42761-021-00089-z

Tingley, D., Yamamoto, T., Hirose, K., Keele, L., & Imai, K. (2014). mediation: R Package for Causal Mediation Analysis. Journal of Statistical Software, 59(5). 10.18637/jss.v059.i05

Trope, Y., Liberman, N., & Wakslak, C. (2007). Construal Levels and Psychological Distance: Effects on Representation, Prediction, Evaluation, and Behavior. Journal of Consumer Psychology, 17(2), 83–95. 10.1016/S1057-7408(07)70013-X

Tsoi, L., Burns, S. M., Falk, E. B., & Tamir, D. I. (2022). The promises and pitfalls of functional magnetic resonance imaging hyperscanning for social interaction research. Social and Personality Psychology Compass, 16(10), e12707. 10.1111/spc3.12707

Tunçgenç, B., & Cohen, E. (2018). Interpersonal movement synchrony facilitates pro-social behavior in children’s peer-play. Developmental Science, 21(1), e12505. 10.1111/desc.12505

Valdesolo, P., & DeSteno, D. (2011). Synchrony and the social tuning of compassion. Emotion, 11(2), 262–266. 10.1037/a0021302

Van Knippenberg, D., De Dreu, C. K. W., & Homan, A. C. (2004). Work Group Diversity and Group Performance: An Integrative Model and Research Agenda. Journal of Applied Psychology, 89(6), 1008–1022. 10.1037/0021-9010.89.6.1008

Vera, D., & Crossan, M. (2005). Improvisation and Innovative Performance in Teams. Organization Science, 16(3), 203–224. 10.1287/orsc.1050.0126

Wallot, S., Mitkidis, P., McGraw, J. J., & Roepstorff, A. (2016). Beyond Synchrony: Joint Action in a Complex Production Task Reveals Beneficial Effects of Decreased Interpersonal Synchrony. PLOS ONE, 11(12), e0168306. 10.1371/journal.pone.0168306

Walton, R. E., & McKersie, R. B. (1965). A behavioral theory of labor relations. McGraw-Hill.

Wang, L.-S., Cheng, J.-T., Hsu, I.-J., Liou, S., Kung, C.-C., Chen, D.-Y., & Weng, M.-H. (2022). Distinct cerebral coherence in task-based fMRI hyperscanning: Cooperation versus competition. Cerebral Cortex, 33(2), 421–433. 10.1093/cercor/bhac075

Weingart, L. R., Brett, J. M., Olekalns, M., & Smith, P. L. (2007). Conflicting social motives in negotiating groups. Journal of Personality and Social Psychology, 93(6), 994–1010. 10.1037/0022-3514.93.6.994

Weingart, L. R., Thompson, L. L., Bazerman, M. H., & Carroll, J. S. (1990). TACTICAL BEHAVIOR AND NEGOTIATION OUTCOMES. International Journal of Conflict Management, 1(1), 7–31. 10.1108/eb022670

Westgate, E. C., & Wilson, T. D. (2018). Boring thoughts and bored minds: The MAC model of boredom and cognitive engagement. Psychological Review, 125(5), 689–713. 10.1037/rev0000097

Wheeler, S. C., Briñol, P., & Hermann, A. D. (2007). Resistance to persuasion as self-regulation: Ego-depletion and its effects on attitude change processes. Journal of Experimental Social Psychology, 43(1), 150–156. 10.1016/j.jesp.2006.01.001

Williams, M., Belkin, L. Y., & Chen, C. C. (2020). Cognitive Flexibility Matters: The Role of Multilevel Positive Affect and Cognitive Flexibility in Shaping Victims’ Cooperative and Uncooperative Behavioral Responses to Trust Violations. Group & Organization Management, 45(2), 181–218. 10.1177/1059601120911224

Wiltermuth, S. S., & Heath, C. (2009). Synchrony and Cooperation. Psychological Science, 20(1), 1–5. 10.1111/j.1467-9280.2008.02253.x

Yeomans, M., Brooks, A. W., Huang, K., Minson, J., & Gino, F. (2019). It helps to ask: The cumulative benefits of asking follow-up questions. Journal of Personality and Social Psychology, 117(6), 1139–1144. 10.1037/pspi0000220

Zhang, H., Yang, J., Ni, J., De Dreu, C. K. W., & Ma, Y. (2023). Leader–follower behavioural coordination and neural synchronization during intergroup conflict. Nature Human Behaviour, 7(12), 2169–2181. 10.1038/s41562-023-01663-0

Zhang, W., Sjoerds, Z., & Hommel, B. (2020). Metacontrol of human creativity: The neurocognitive mechanisms of convergent and divergent thinking. NeuroImage, 210, 116572. 10.1016/j.neuroimage.2020.116572

